# Role of nanoscale antigen organization on B-cell activation probed using DNA origami

**DOI:** 10.1101/2020.02.16.951475

**Authors:** Rémi Veneziano, Tyson J. Moyer, Matthew B. Stone, Tyson R. Shepherd, William R. Schief, Darrell J. Irvine, Mark Bathe

## Abstract

Arraying vaccine immunogens in a multivalent form on the surface of virus-like particles is an important strategy used to enhance the efficacy of subunit vaccines. However, the impacts of antigen valency, spacing, and spatial organization on B cell triggering remain poorly understood. Here, we use DNA origami nanoparticles to create precise nanoscale organizations of a clinically-relevant HIV gp120 immunogen to systematically interrogate their impact on B cell triggering *in vitro*. We find that antigen dimers elicit monotonically increasing B cell receptor activation as inter-antigen spacing increases up to ~30 nm, and only 5 immunogens arrayed on the surface of a 3D particle are needed to elicit maximal B cell calcium signaling. Our results reveal design principles of viral and immunogen display that drive functional B cell responses.

## MAIN TEXT

Efficient activation of antigen-specific B cells is a central goal in the design of new vaccines. One well established strategy to enhance B cell activation employs multivalent presentation of immunogens^1^. Antigen multimers, antigen-conjugated polymers, and viral-like nanoparticles (NPs) displaying immunogens at high density have all been shown to effectively crosslink B cell receptors (BCRs) and strongly initiate early B cell signaling^2–6^. Antigen multimerization enables very low affinity B cells to be fully activated^7,8^, which may be particularly important for approaches such as lineage-guided or germline-targeting vaccines that aim to activate specific rare, low-affinity target B cell populations. However, the roles of distinct antigen spatial arrangements on triggering B cells and initiating robust BCR signaling remain poorly understood. To date, studies exploring the effect of antigen organization on B cell triggering have generally employed protein, polymer, or particle scaffolds that only allowed limited variation of spatial parameters, or provided only statistical control over the numbers and locations of antigens^2,3,9–12^.

In order to independently probe the relative roles of immunogen valency and spacing on BCR activation, here we used scaffolded DNA origami NPs^13,14^ to display discrete antigen copy numbers with controlled inter-antigen spacings on the scale of an individual virus-like NP^15,16^. As a model antigen, we employed an HIV germline-targeting gp120 protein, the engineered outer domain eOD-GT8. This immunogen is designed to bind with high affinity to the unmutated common ancestor of the VRC01 CD4 binding site-specific HIV broadly neutralizing antibody (glVRC01), and thereby activate a collection of human naïve B cells expressing so-called VRC01-class BCRs^17–19^. eOD-GT8 activates both cognate glVRC01 BCR-expressing cell lines and murine BCR-transgenic primary B cells, but only when presented to B cells in multivalent form^8,17^. We previously showed that multimerization of 60 copies of eOD-GT8 via fusion to the self-assembling bacterial protein lumazine synthase formed an isotropic ~30 nm diameter NP (eOD-60mer) that elicited robust B cell activation *in vitro* and *in vivo*^17,20,21^.

Motivated by these findings, we designed two types of structured DNA-NPs^13,14^ to interrogate the specific roles of antigen copy number and spatial patterning on B cell triggering: A 3D icosahedral NP^14^ with a ~40 nm diameter size comparable to the eOD-60mer, and a 80 nm long rigid-rod six-helix bundle (6HB) (**Fig. 1a and Supplementary figs. 1 and 2**). eOD-GT8 antigens were site-specifically coupled to these nanostructures through hybridization to free single-stranded DNA (ssDNA) overhang strands displayed from the origami at specific programmed locations (**Supplementary fig 3**). Using this approach, we were able to present the antigen in copy numbers varying from 1 to 60 in variable nanoscale spatial organizations, enabling independent control over antigen stoichiometry, inter-antigen distance, as well as the spatial dimensionality of antigen presentation (**Fig. 1b**). As comparative controls to probe the roles of rigidity and distance on BCR signaling, we also prepared dimer constructs based on flexible ssDNA or polyethylene glycol (PEG) polymer linkers of varying lengths (**Supplementary Table 1**).

**Fig. 1.**
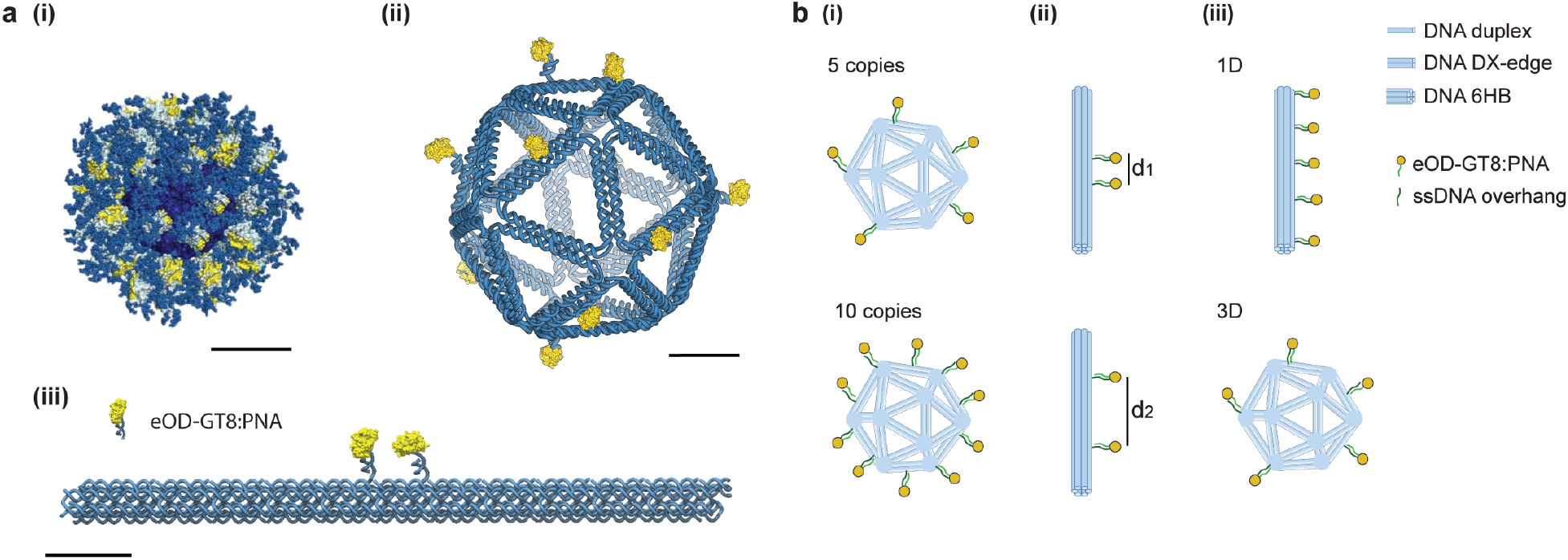
Scaffolded DNA origami NPs to control nanoscale organization of HIV immunogens. **(a)** DNA-NPs were designed to self-assemble the eOD-GT8 antigen in a controlled manner, mimicking features of the eOD-GT8-60mer immunogen. (**i**) eOD-GT8-60mer protein NP; (**ii**) Icosahedral DNA-NP presenting 10 copies of eOD-GT8 (Ico-10x); and (**iii**) 6HB rod-like structure presenting two copies of eOD-GT8 (6HB-2x). Scale bars are 10 nm. **(b)** Both the icosahedral and 6HB structures were used to explore the **(i)** stoichiometry; **(ii)** inter-antigen distance, d; and **(iii)** 1D versus 3D dimensionality of presentation of eOD-GT8 antigens.

Negative stain transmission electron microscopy (TEM) imaging and agarose gel electrophoresis of the 6HB and icosahedral DNA-NP constructs confirmed their geometry, monodispersity, and structural rigidity, consistent with previous work^14^ (**Fig. 2a and Supplementary figs. 4-7**). Short, outward-facing ssDNA overhangs were used at the 3’ ends of select DNA-NP staple strands to anchor individual eOD-GT8 monomers (**Supplementary fig. 3**) via hybridization to a complementary peptide nucleic acid tag site-specifically conjugated to the antigen (**Supplementary figs. 8-10**). eOD-GT8-PNA monomers were purified and added to pre-folded and purified DNA-NPs to allow for complete hybridization prior to purification and application to B-cells *in vitro*. Antigen-coupled NPs showed expected shifts in gel electrophoresis (**Fig. 2b**), while fluorimetry measurements (using a fluorescent version of the PNA-eOD-GT8 labeled with AlexaFluor647) and Tryptophan fluorescence measurement confirmed efficient and stable complexation of eOD-GT8-PNA to DNA-NPs with quantitatively controlled stoichiometries varying from 2 to 60 (**Fig. 2b, Supplementary figs. 11-13, and Supplementary Table 2**).

**Fig. 2.**
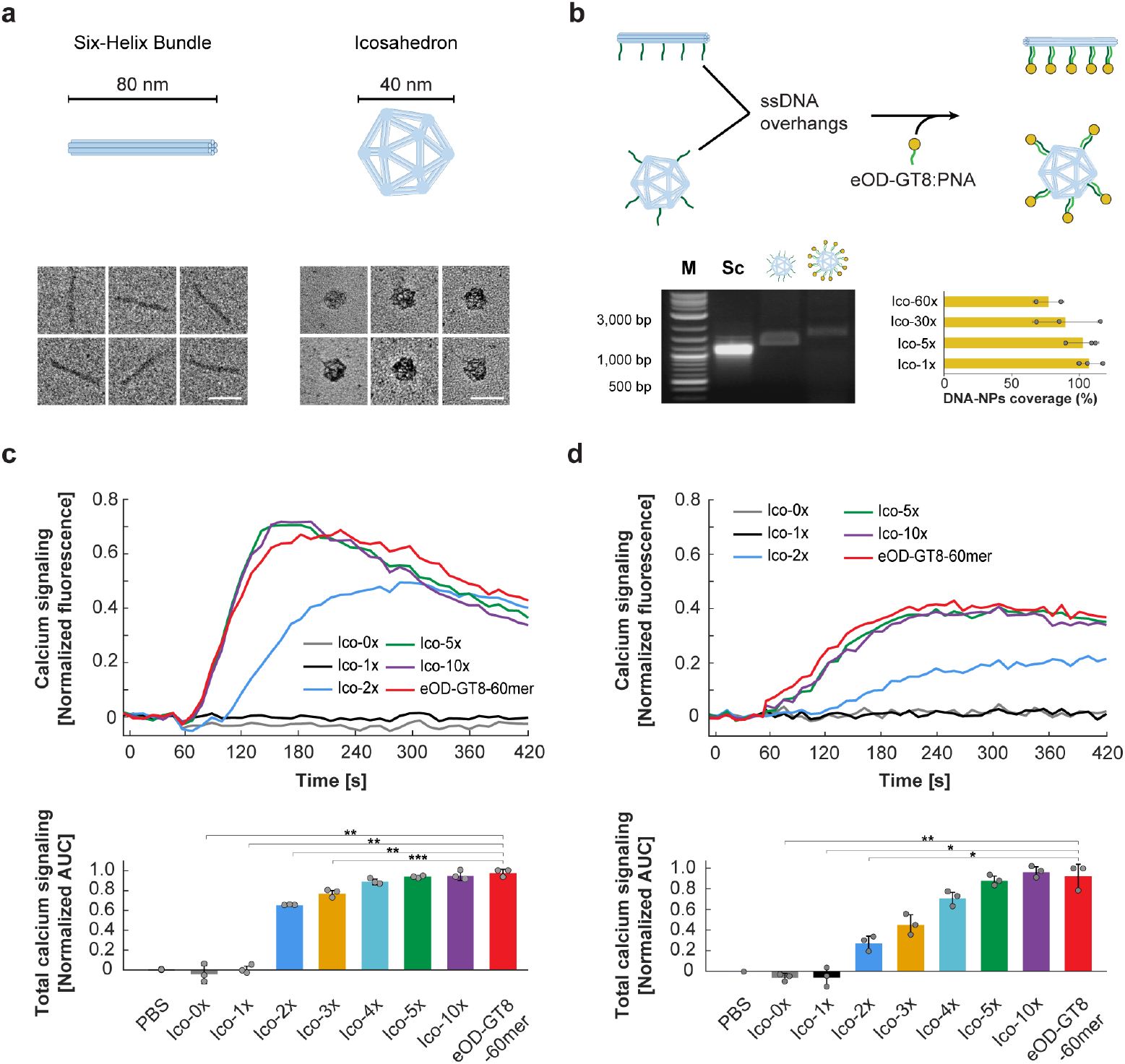
Increasing antigen valency improves B cell responses to NP antigens up to a threshold. **(a)** Folding of the two types of DNA-NPs (six-helix bundle, 6HB, and DNA icosahedron) that were designed and used in this study for 1D versus 3D presentation of antigens. TEM images show high folding yield and monodisperse DNA-NPs. **(b)** Overview of the antigen conjugation protocol to attach eOD-GT8 antigens to the DNA-NPs using PNA single strands complementary to ssDNA overhangs on the DNA-NPs and characterization with electrophoresis. Shown are representative gel electrophoresis samples and fluorescence quantification of four different icosahedral DNA-NPs conjugated with eOD-GT8:PNA-AF647 (Ico-1x; Ico-5x; Ico-30x; Ico-60x)); (L, molecular weight ladder; Sc, Scaffold). Error bar represents standard deviation of the mean (n= 3 samples/group for Ico-1x, Ico-5x, Ico-30x; n=2 samples/group for Ico-60x). **(C and D)** DNA-NPs modified with eOD-GT8 activate BCR at both **(c)** 5 nM and **(d)** 0.5 nM eOD-GT8. Fluo-4 calcium probe fluorescence is shown (representative individual calcium traces). in the top row, and average areas under the time-course are normalized to the maximum response of all samples in a repeat shown in the bottom row. Error bars represent standard deviation of the mean (n=3 samples/group). P-values are from Student’s t-test (*: p<0.05; **: p<0.01; ***: p<0.001).

We first analyzed the impact of eOD-GT8 valency on B cell responses using a ~40 nm diameter icosahedral DNA-NP bearing zero to ten copies of eOD-GT8 monomers distributed equidistantly from one another over the surface of the NP (**[0-10]-mer, Supplementary fig. 14 and Supplementary Table 3**). As a readout of BCR triggering, we focused on dynamic measurements of intracellular calcium signaling as a critical signature of full BCR activation^22,23^. Human Ramos B cells stably expressing the cognate germline VRC01 IgM receptor were incubated with DNA-NPs equivalent to 0.5 nM or 5 nM total eOD-GT8, or the same eOD-equivalent concentration of eOD-60mer protein NPs, and B cell responses were recorded spectroscopically using a fluorescent intracellular calcium indicator dye. As expected^8,17^, antigen-free DNA-NPs or NPs bearing a single copy of eOD failed to activate B cells at either antigen concentration. By contrast, NPs bearing two or more copies of eOD-GT8 triggered monotonically increasing cellular responses with increasing antigen valency (**Fig. 2c and d**). At both antigen concentrations, signaling plateaued in response to 5-mer or higher valency DNA-NPs, at a level indistinguishable from the eOD-GT8-60mer. Moreover, increasing the valency on the DNA icosahedron to 30 or 60 copies of eOD-GT8 had no further effect on increasing cellular response (**Supplementary fig. 15)**. Because the affinity of eOD-GT8 for glVRC01 is high, with a K_D_ ~ 30 pM^17^, we also tested the same icosahedral DNA-NP functionalized with a lower-affinity variant of the gp120, eOD-GT5 (K_D_ ~0.5 μM [^8^]) **(Supplementary fig. 16**). As previously observed with eOD-GT8, the five and ten 10 copy number NPs exhibited a similar plateau in signaling to the icosahedral constructs conjugated with eOD-GT5. Importantly, we noted that even at an eOD-GT5 concentration of 25 nM, the maximum activation remained lower than that observed using the eOD-GT8-60-mer, and using the eOD-GT8 DNA icosahedron and the eOD-GT8 60-mer at lower immunogen concentrations (2 nM).

We next sought to define the impact of antigen spacing on BCR triggering. Previous studies have shown that inter-antigen distances impact receptor signaling for both Fc receptor engaging IgE^24^ and the T cell receptor^25^, and that the oligomeric organization of the BCR impacts receptor activation^26^. To evaluate antigen spacing in the absence of additional 3D spatial variables, we turned to the 6HB rigid rod origami as a platform to present antigen with precise 1D spacings (**Supplementary figs. 2 and 3**). Unexpectedly, calcium signaling in responding B cells began at a minimum spacing of 7 nm (+/− 3 nm linker size), and subsequently increased monotonically with increasing antigen spacing at fixed total antigen concentration (**Fig. 3a**). Further, calcium response plateaued at an antigen spacing of 28 nm, yet was still maximal at a spacing of 80 nm despite the fact that this spacing would definitively preclude close approach of two BCRs bound to the same rod. Enhanced triggering by the more widely-spaced dimers was not due to differences in overall engagement with the nanostructures because flow cytometry analysis of fluorescent 6HB association with Ramos cells showed identical binding of 7 nm-spaced eOD dimers and 28 nm-spaced dimers to B cells after a 30-minute incubation with the constructs (**Fig. 3b and Supplementary fig. 17**).

**Fig. 3.**
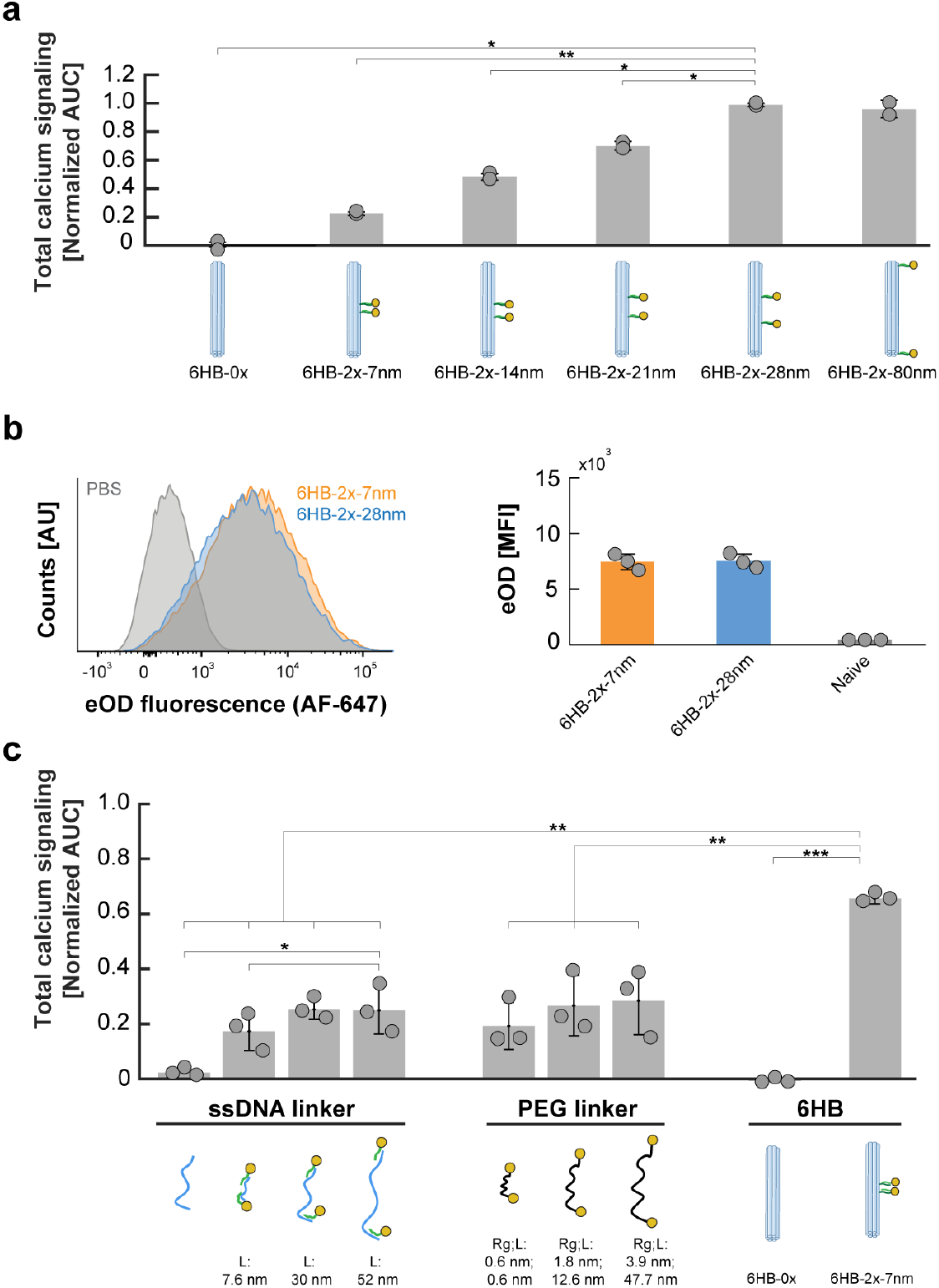
BCR response increases and then plateaus with increasing inter-antigen distance on a rigid scaffold. **(a)** Area-under-the-curve total calcium signaling in glVRC01 B cells stimulated with DNA-NP eOD-GT8 dimers with inter-antigen distances between 7 nm and 80 nm at an antigen concentration of 5 nM (n=2 samples/group). **(b)** Representative flow cytometry plots of 6H-2x-7nm, and 6HB-2x-28nm binding to antigen-specific B cells (left) after 30 min incubation at 4°C. (Right) Quantitation of data from flow cytometry (MFI: Mean Fluorescence Intensity). Error bars represent standard deviation (n=3 samples/group). **(c)** Total calcium release (Fluo-4 fluorescence) integrated over 7 minutes following antigen addition from cells stimulated with eOD-GT8 dimers attached to the flexible polymeric scaffolds (ssDNA or PEG), compared with rigid 6HB DNA-NP eOD-GT8 dimer structures at a fixed antigen concentration of 5 nM (n=3 samples/group). Fluo-4 AUC is normalized as in Figure 2, where error bars represent standard deviations (n=3 samples/group) and P-values are from Student’s t-test (*: p<0.05; **: p<0.01; ***: p<0.001). (L: Contour Length; Rg: Radius-of-Gyration).

To probe the role of scaffold rigidity in BCR triggering by dimeric antigens, we compared calcium signaling induced by DNA rods to eOD-GT8 dimers presented using flexible ssDNA or PEG linkers of variable contour lengths (**Supplementary Table 1**). These flexible polymer constructs presenting eOD-GT8 dimers elicited substantially reduced B cell signaling compared with their rigid DNA-NP counterparts, indicating the importance of both linear distance and also rigidity of the structural scaffold itself for inducing robust B cell response (**Fig. 3c**). These results are of interest in light of past studies demonstrating the importance of B cell response to the mechanical properties of antigen presenting cells^5,27^.

The preceding dimer results suggest that clustering of antigens below a threshold distance may in fact limit BCR responses, in contrast to a model where tight clustering of antigen is needed for maximal BCR response by facilitating inter-BCR cooperativity^28–30^. To further elucidate the roles of both inter-antigen spacing and dimensionality of presentation on B cell response, we programmed different clusters of five eOD-GT8 antigens on one face of the icosahedral DNA-NP using distinct inter-antigen distances of ~3 to 22 nm (**Fig. 4a and b**). We found that increasing inter-antigen distances on the DNA icosahedron led to an increase in the cellular response similar to what was observed with the dimer results presented in Figure 3. Using the linear rods to cluster 5 antigens with similar inter-antigen distances shows that decreasing the distance between the antigens in both the planar and the linear presentations yielded decreasing cellular responses, consistent with observations using the 6HB dimer DNA-NP (**Supplementary fig. 18**). However, for smaller inter-antigen distances (namely 7 and 11 nm), the linear antigen placement yielded a greater response than the clustered, planar placement of antigens (**Supplementary fig. 18**), suggesting that the maximum spacing (rather than the minimum spacing) between any two antigens in the array determines the observed level of activation.

To probe the mechanism of B cell activation, we examined three distinct DNA-NP constructs using fluorescence imaging: two 6HB dimers with inter-antigen spacings of 7 and 28 nm that respectively triggered low and high B cell activation, and one DNA icosahedral NP presenting 30 copies of eOD-GT8 that triggered robust B cell activation. Fluorescently labeled eOD-GT8-presenting DNA-NPs were added to Ramos cells at identical total eOD-GT8 concentrations, and spatially correlated with fluorescently labeled BCR on the cell surface prior to internalization (**Fig. 5a and Supplementary figs. 19-20**). As anticipated, DNA-NP binding was strongly correlated with VRC01 IgM expression, whereas cells lacking IgM expression failed to bind the eOD-GT8-bearing particles (**Fig. 5b and Supplementary figs. 19-20**). For all DNA-NP constructs examined, antibody staining of phosphorylated Syk kinase (pSyk) revealed a sharp increase in pSyk phosphorylation after only 1 minute of DNA-NP or eOD-GT8-60mer addition. Moreover, dimers of eOD-GT8 separated by short distances of only 7 nm (6HB-2x-7nm) resulted in significantly lower pSyk accumulation compared with dimers separated by larger distances of 28 nm (6HB-2x-28nm) or icosahedral NPs bearing 30 copies of eOD-GT8 (Ico-30x) (**Fig. 5c**). Characterization of the internalization of eOD using phalloidin to stain filamentous actin in the actin-rich cellular cortex indicated greater internalization of the icosahedral 30-mer NP compared with its 6HB-2x-7nm and 6HB-2x-28nm counterparts (**Fig. 5d**). Taken together, these results corroborated both the levels of cellular activation observed for these three DNA-NP constructs and indicate that the previously observed calcium responses were acting through the BCR-mediated activation pathway that includes pSyk phosphorylation and downstream NP internalization.

**Fig. 4.**
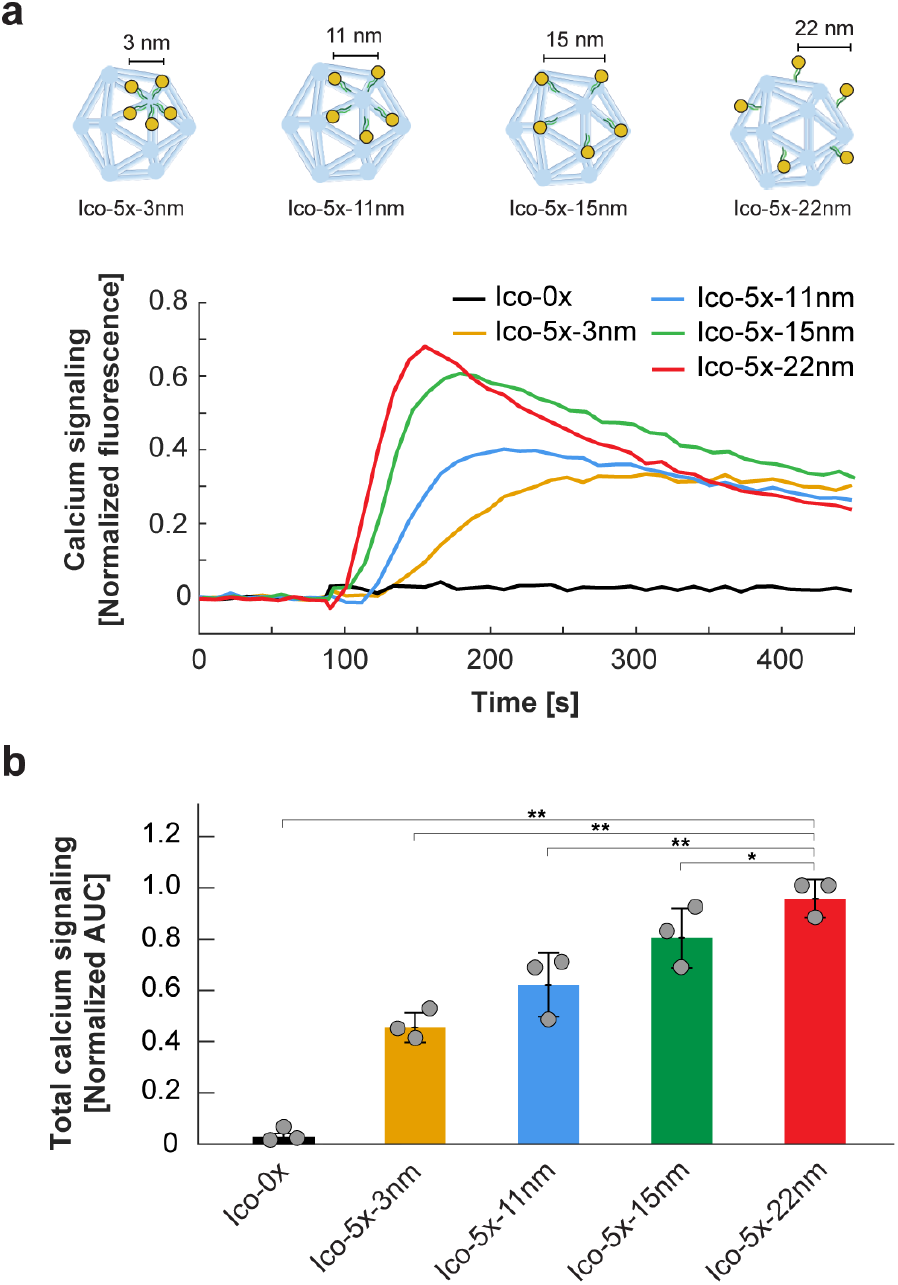
Clustering of antigens on one face of an icosahedral DNA-NP. **(a)** Fluo-4 calcium probe fluorescence versus time following addition of 5 nM eOD-GT8 antigen to glVRC01 B cells. Icosahedral (Ico) structures with varying inter-antigen distances are plotted in the same graph (representative individual calcium traces). (**b)** Total calcium signaling integrated over 6 min following antigen addition from cells stimulated with different Ico-5x structures presenting antigen at a total concentration of 5 nM eOD-GT8 (n=3 samples/group). Fluo-4 AUC is normalized as in Figure 2, where error bars represent standard deviations and P-values are from Student’s t-test (*: p<0.05; **: p<0.01; ***: p<0.001).

**Fig. 5.**
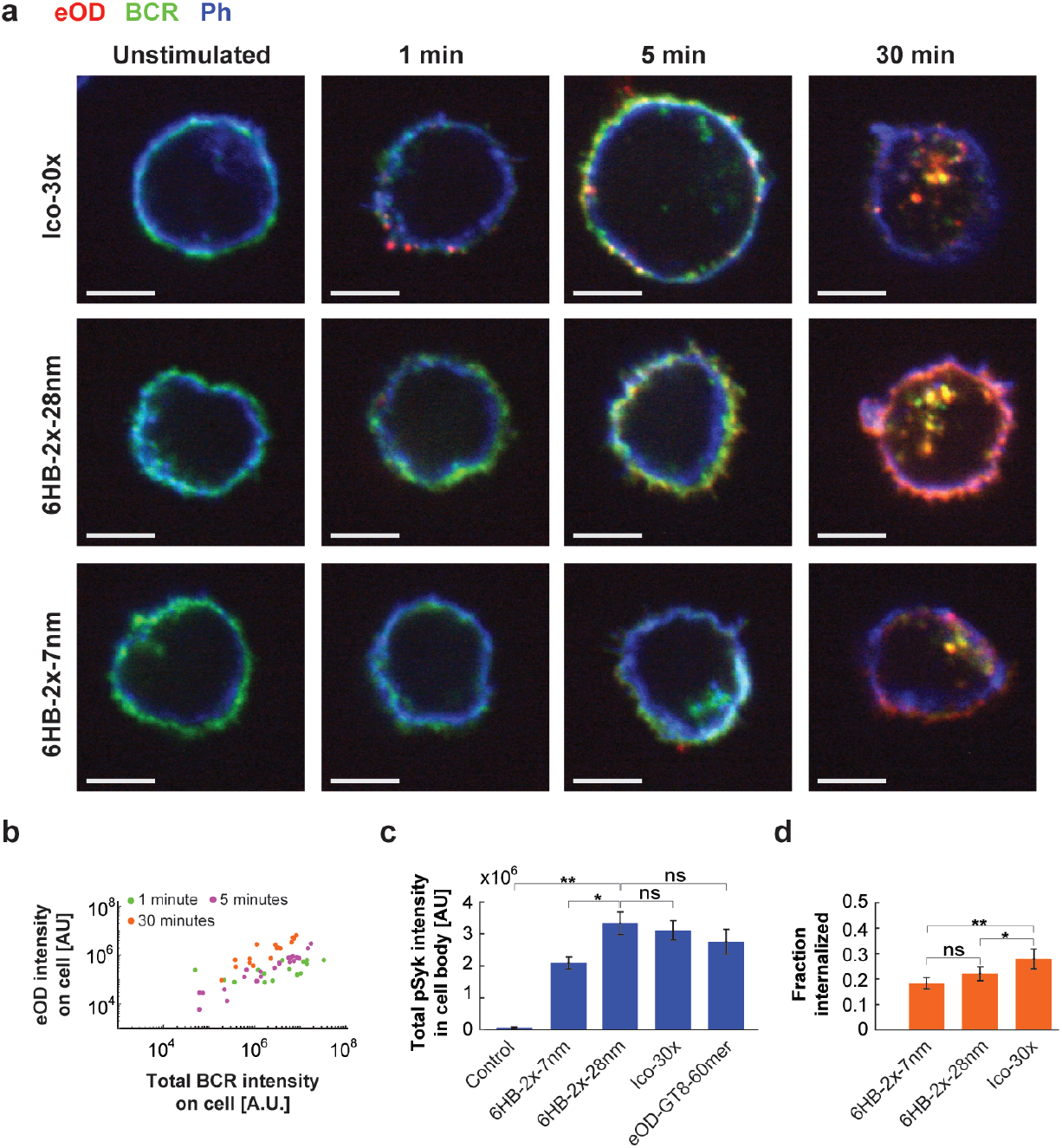
Confocal microscopic imaging of DNA-NPs on Ramos B cells. **(a)** Time series imaging of Ico-30x, 6HB-2x-28nm, and 6HB-2x-7nm shows surface binding and internalization into Ramos B-cells. Here, eOD-GT8 was fluorescently labelled with Alexa Fluor 647, VRC01 IgM BCR was labeled with a FAb fragment conjugated to Janelia Fluor 549 prior to antigen addition, and actin was labeled with phalloidin Alexa Fluor 405 after cell fixation. Scale bar is 5 μm. **(b)** Total intensity of eOD-GT8 is highly correlated with intensity of BCR, confirming specific binding of NPs to the BCR and co-internalization. **(c)** Ramos cells were labeled with an anti-phospho-Syk antibody after fixation and the total pSyk intensity per cell was determined. Numbers of cells analyzed are 11 (control), 58 (6HB-2x-7nm), 43 (6HB-2x-28nm), 56 (Ico-30x), and 56 (eOD-GT8-60mer). **(d)** Internalized fraction of eOD was estimated by segmenting the cell surface using a phalloidin stain as detailed in Methods. Total internal eOD fluorescence was divided by total cellular eOD fluorescence on a cell-by-cell basis. Numbers of cells analyzed are 19 (6HB-2x-7nm), 23 (6HB-2x-28nm), and 15 (Ico-30x). In both C and D, error bars denote the standard error of the mean fluorescence between cells with significance determined by Student’s t-test (*: p<0.05; **: p<0.01; ***: p<0.001).

We have observed that distances between immunogens are crucial determinants of BCR activation and cellular response. Overall, our data are consistent with previous observations that antigen multivalency is a necessary feature for efficient triggering of BCRs. However, in contrast to a model in which tight antigen clustering is needed to yield greater B cell activation, here we find that extended antigen separation on rigid scaffolds leads to an equivalent and in some cases superior BCR activation. Intriguingly, both linear and planar antigen presentations yielded similar cellular responses when the inter-antigen distance was greater than a critical threshold of ~15 nm, suggesting that the average distances and the dimensionality between the antigenic sites does not drive the observed differences between the constructs above this critical distance. Moreover, our results showing efficient signaling induction for an 80 nm dimer (**Fig. 3a**) suggest that non-local interactions between antigen receptors may drive the initiation of signaling in response to antigen binding. These non-local interactions may stem from signaling cooperativity between the antigen receptor and slowly evolving structural elements in the B cell membrane^31^ and the actin cortex^32^, which could facilitate communication of receptor binding over such large distances.

We hypothesize that the observed decrease in signaling response of B cells to dimers with short inter-antigen distances is due to a bivalency effect, whereby single immunoglobulins are binding to both eOD sites when inter-antigen spacing is below ~28 nm, thereby resulting in engagement of only single BCRs. This hypothesis would further underline the importance of coordination between multiple antigen receptors for efficient receptor signaling, as shown previously using programmed DNA assemblies^15,16^. Affinity is an important determination of BCR responses^33,34^, where it is important to note here that the affinity between germ-line VRCO1 and eOD-GT8 is in the sub-nanomolar range, which places this system in the regime of mature BCR antigen affinities rather than naïve affinities. In the case of low affinity interactions, excessive multivalency may play a role in improving the avidity between a virus and B cell. It has been suggested that the low number of Env viral spikes displayed on the viral surface may help viruses to evade the immune system^35^. In light of our findings, this low spike density might impact the overall avidity between HIV and the immature B cell surface, while having minimal impact on the induction of mature, high affinity receptor signaling during viral encounters. Importantly, the rigid nature of DNA origami nanostructures provides an effective scaffold for precisely defining spatial relationships between immuongens, and as we show here, can be used to define design rules relevant for protein-based virus-like NPs that can be computationally designed to present rigid immunogen arrays following these criteria^36^.

## ACKNOWLEDGEMENTS

This work was supported by the Human Frontier Science Program (RGP0029/2014), the Office of Naval Research (N00014-16-1-2953), the U. S. Army Research Office through the Institute for Soldier Nanotechnologies at MIT (Cooperative Agreement Number W911NF-18-2-0048), the Ragon Institute of MGH, MIT, and Harvard, and the NIH (R21-EB026008, R01-MH112694, AI048240, and UM1AI100663). DJI is an investigator of the Howard Hughes Medical Institute.

## AUTHOR CONTRIBUTIONS

R.V. designed the different DNA-NPs and performed the folding and characterization of the DNA-NPs, the conjugations with different antigens, and their fluorescence quantification measurements, and analyzed the data. T.J.M. performed the antigen modification with PNA and fluorescent dye, the B cell calcium flux assay, the flow cytometry, and analyzed the data. M.B.S. performed the confocal microscopy imaging, the image analysis, and analyzed the data. T.R.S. performed the TEM characterization. W.R.S. designed the immunogens. M.B. and D.J.I. designed and supervised the study and interpreted the results. R.V., T.J.M., M.B.S., D.J.I., and M.B. wrote the manuscript. All authors commented on and edited the manuscript.

## METHODS

### Chemicals and kits

Magnesium chloride, TRIS acetate EDTA (TAE) buffer, TRIS-base, sodium chloride, Phosphate Buffer Saline, ethidium Bromide solution (10 mg/mL) and Amicon ultra 0.5 centrifugal filter (#UFC5003) were provided by Sigma-Aldrich. Nuclease free water was purchased from Integrated DNA Technologies, Inc. (IDT). The DNTPs mix (#N0447S), the DNA ladder (Quick-Load^®^ Purple 2-Log DNA ladder 0.1-10 kb, #N0550S) were provided by New England Biolabs (NEB), The polymerase enzyme (Accustart Taq DNA polymerase HiFi, #95085-05K) was provided by Quanta Biosciences. Low melt agarose was purchased from IBI Scientific (#IB70058) and the agarose from Seakem, Inc. G-capsule for electroelution (#786-001) was purchased from G-Biosciences and Freeze ‘N Squeeze DNA gel extraction columns by Bio-rad, Inc. (#732-6165). The Zymoclean Gel DNA recovery kit (#D4008) was purchased from Zymo Research, Inc. The SybrSafe DNA staining reagent was provided by Thermo Fisher Scientific, Inc. PEG3500 (#A4010-1/MAL-PEG3500-MAL) and PEG2000 (#A4010-1/MAL-PEG2000-MAL) bismaleimide were purchased from JenKem Technology.

### Oligonucleotides and DNA templates

All oligonucleotides used for asymmetric PCR (aPCR) amplification of the template and for folding the various scaffolded DNA origami nanoparticles (NPs) were purchased from IDT. The circular plasmid DNA scaffold M13mp18 used for amplification of the short scaffolds with aPCR (see Supplementary sequences document) was acquired from NEB (#N4040S).

### Antigens and cell lines

The eOD antigen with an N-terminal cysteine was produced in HEK cells, and purified by affinity chromatography using a Nickel affinity column. The protein was then further purified by size exclusion chromatography using a Superdex 75 10/300 column (GE Healthcare). Ramos B cells expressing VRC01 germline B cell receptor were provided by Daniel Lingwood (Ragon Institute).

### ssDNA scaffold synthesis

The ssDNA scaffolds used to fold the DNA six helix bundle (6-HB) and the DNA icosahedron NPs were produced using the previously described procedure asymmetric PCR (1, 2). Briefly, two specific primers sets were used to amplify the ssDNA fragments (supplementary Table S1) using Quanta Accustart HiFi DNA polymerase. The aPCR mix was prepared at a final volume of 50 μL with the specific polymerase buffer complemented with 2 mM magnesium chloride, 200 μM dNTPs, 1μM forward primer, 20 nM reverse primer, 25 ng M13mp18 template, and 1 unit of Quanta Accustart HiFi polymerase. The amplification protocol used was: 94°C for 1 min for initial denaturation followed by 35 cycles of 94°C held for 20 sec; 56°C held for 30 sec; 68°C held for 1 min per kb for amplification. Following amplification, the aPCR mix was run on a 1% low-melt agarose gel prestained with Ethidium Bromide (EtBr). The resulting ssDNA product was then extracted using the Zymoclean gel DNA recovery kit. Purified ssDNA concentration was measured using a NanoDrop 2000.

### DNA-NP folding

DNA-NPs (Icosahedron and 6-HB) with or without overhangs were self-assembled using a one-pot reaction and annealing as described previously^13,37^. Briefly, 20-40 nM of scaffold was mixed with an excess of the correct staple strand mix (molar ratio of 10x) in buffer TAE-MgCl_2_ (40 mM Tris, 20 mM acetic acid, 2 mM EDTA, 16 mM MgCl_2_, pH 8.0) in a final reaction volume of 50 uL and annealed with the following program: 95°C for 5 min, 80–75°C at 1°C per 5 min, 75–30°C at 1°C per 15 min, and 30–25°C at 1°C per 10 min. The folded NPs are stored at 4°C in the folding buffer with the excess of staples strands prior to perform conjugation with antigens.

### DNA-NP purification

Before using the DNA-NPs for conjugation with antigens and for the B cell activation assay, the DNA origami objects folded with an excess of staples strands were purified using an Amicon ultra 0.5 centrifugal filter with three washes of folding buffer and an extra wash of 1X PBS for further modification with antigens. Centrifugation steps were performed at 1000g for 30-40 minutes and the final concentration of NPs was determined using a NanoDrop 2000. Purified NPs were subsequently modified with antigens or stored in 1X PBS at 4°C.

### PNA strand synthesis

PNA strands were synthesized in-house using solid phase peptide synthesis. Lysine residues were attached at either end of the PNA sequence to improve solubility. Fmoc-PNA monomers (PNA-Bio, Inc.) were coupled to a low loading Tentagel-S-RAM resin using 4 eq. PNA, 3.95 eq. PyBOP, and 6 eq. diisopropylethylamine (DIEA). Lysine and glycine residues were reacted in the same way. Following each coupling, the peptide was deprotected in 20% piperidine in DMF. N-maleoyl-β-alanine (Sigma) was coupled to the N-terminus under the same coupling conditions.

The peptide was then cleaved from the resin in 95% trifluoroacetic acid (TFA), 2.5% H2O, and 2.5% triisopropylsilane. The peptide was dissolved in aqueous solution with 0.1% TFA, filtered, and purified by HPLC using a C-18 Gemini column (Phenomenex) with a mobile phase of acetonitrile containing 0.1% TFA. Purity of the PNA products was analyzed with MALDI-TOF mass spectrometry on a Bruker Daltonics microflex. The sequence of the synthesized PNA strand is: (Maleimide)-GGK-cagtccagt-K-(CONH_2_), and the complementary ssDNA is: 5’-Oligo-TT-ACTGGACTG-3’ (melting temperature predicted: 56.7°C). The sequence has been designed to have a melting temperature above 55°C (predicted with the PNA tool: https://www.pnabio.com/support/PNA_Tool.htm, from PNA Bio, Inc.) and orthogonal to the sequence of M13mp18 and validated using NCBI BLAST online tool^38^.

### Antigen-PNA conjugation

PNA strands were conjugated to eOD by reacting the terminal maleimide onto an N-terminal cysteine of eOD. Prior to the reaction, eOD was incubated with a 10-fold molar excess of tris(2-carboxyethyl)phosphine (TCEP) for 15 minutes, after which TCEP was removed using a centrifugal filter. Immediately after removal of TCEP, a 2-fold molar excess of maleimide-PNA was reacted with cysteine-eOD overnight at 4C in PBS. Unreacted PNA was then removed using an Amicon centrifugal filter (10 kDa MWCO).

### Antigen conjugation with AF647 dye

The eOD-PNA conjugate was modified with the fluorescent label AlexaFluor 647-NHS (AF647). eOD-PNA was incubated with 5 molar equivalents of AF647-NHS in 10 mM sodium bicarbonate buffer for 2 hours at room temperature. Unreacted dye was removed using centrifugal filtration (10kDa MWCO).

### Antigen attachment to DNA-NPs

Purified DNA-NPs were mixed with PNA-antigen conjugates at a molar ratio of 5X antigen per overhang on the DNA-NPs in 1X PBS buffer. The concentration of DNA-NPs used was in the range of 50 to 100 nM. An annealing temperature ramp was used for ssPNA-ssDNA hybridization starting at 37°C and decreasing to 4°C at 1°C per 20 min and kept for at least 4 hours at 4°C prior use for B cell activation assay.

### Structural characterization

#### Transmission Electron Microscopy

DNA-NPs were visualized by transmission electron microscopy (TEM) using grids prepared as described previously with minor modifications^39^. Briefly, carbon supported grids with copper mesh (CF200H-CU; Electron Microscopy Sciences) were glow discharged and soaked in 100 μM MgCl2 and blotted prior to depositing DNA-NPs. 20 μl of a 10 nM DNA-NP solution was applied to a clean parafilm surface and the grid was floated for 2 minutes. While soaking, 2% uranyl formate (UF; Electron Microscopy Sciences) was neutralized with 25 mM NaOH final concentration, vortexed for 1 minute, and filtered via syringe through a 0.1 μm filter (EMD Millipore) dropwise onto the clean parafilm surface. The grid was then removed and quickly dried by edge blotting with Whatman 44 ashless paper. The grid was then immediately transferred to the 2% UF solution and incubated for 30 seconds. Again, the grid was dried by blotting along the edge with Whatman paper, and left to dry in air for an additional 30 minutes prior to imaging. Imaging was done on a FEI Tecnai G2 Spirit TWIN set to 120kV equipped with a Gatan camera. Images were acquired at 6,500x for wide-field views and 52,000x for near-field views. Images were collected using 3-second exposures. All raw images were cropped in Adobe Photoshop with subsequent autocontrast applied.

#### Agarose Gel Electrophoresis

DNA-NPs folded and conjugated with eOD-GT8-PNA were analyzed using agarose gel electrophoresis with 2% agarose gel pre-stained with EtBr. Samples non purified in folding buffer or purified in PBS buffer were loaded at a concentration of 20 to 50 nM of DNA origami, ran for 2-3 hours at 70 V at 4C and visualized with a transilluminator. For fluorescence gel analysis with the AF647 modified eOD-GT8 images were acquired using a Typhoon FLA 7000 at the SybrSafe excitation wavelength (473 nm), and at the AF647 excitation wavelength (635 nm). Images were subsequently merged using ImageJ software^40^.

#### Fluorescence quantification

Quantification of the eOD-GT8 conjugation to DNA-NPs was performed using a Fluoromax-4 (Horiba, Inc.) fluorimeter. eOD-GT8-PNA monomers were modified with AF647 dye using NHS-NH_2_ chemistry as described above, prior to conjugation via hybridization to DNA-NPs. eOD-GT8-PNA was incubated with 5 molar equivalents of AF647 for 2 hours, and subsequently purified using centrifugal spin filtration (10k MWCO). The degree of labeling was 2 dyes per protein on average. Spectra were acquired with an excitation wavelength of 630 nm (emission measured at 670 nm). A fluorescence calibration curve was first measured using free monomeric eOD-GT8-PNA conjugated with AF647 dye by varying antigen concentration, and subsequently used as a reference curve to determine the conjugation yield to DNA-NPs.

### B cell Calcium flux assay

Ramos B Cells at a concentration of 10 million cells/mL were incubated with 10 μM Fluo-4 AM (ThermoFisher, Inc.) for 30 minutes at 37C. After washing once, flux assays were performed on a Tecan plate reader at 37C on a 96 well microplate with 160 μL of Fluo-4 labeled Ramos cells at 2 million cells/mL. A baseline fluorescence was then recorded for 1 minute, and 40 μL of NPs were added to the cells for a final concentration of 5 nM of antigen, unless otherwise stated.

### B cell Calcium flux data statistical analysis

Raw calcium traces were normalized to a common baseline by subtracting the PBS timetrace at every timepoint, then dividing the timetrace at every point by the average of the timepoints before antigen addition. The timepoints after antigen addition were then summed for each sample in each repeat to give the calcium release above baseline (I_tot_). The maximum I_tot_ across all samples within each repeat was determined (max(I_tot_)), and total calcium signaling (Normalized AUC) for each sample within each repeat is then given by I_tot_ / max(I_tot_). Repeats were averaged together to yield the bar height for graphs in Figures 2–4. Student’s t-test was performed on the Normalized AUCs entering into this average, where in most cases N–3 replicates.

### B cell imaging

#### Sample preparation for confocal microscopy

Ramos cells were labeled on ice at a concentration of 5 million/mL and protected from light for 30 minutes in Hank’s Buffered Sterile Saline (HBSS) with 20μg/mL human anti IgM f(Ab)_1_ fragment (Jackson 109-007-043) conjugated to Janelia Fluor 549. Cells were spun down and resuspended in warm HBSS at a concentration of 2 million/mL. Antigens were added to a final concentration of 5 nM by adding 50 μL antigen solution to a volume of cells between 175 μL and 400 μL, and cells were kept at 37C by incubation in a thermal bead bath. At timepoints following the addition of antigen, 100 μL of cells were removed and placed into 200 μL of 6% warm PFA solution and allowed to fix for 10 minutes at 37C. Following fixation, fixed cells were diluted in 4.5 mL HBSS and centrifuged at 600g for 5 minutes. Cells were then labeled for 5 hours at 4C in 50 μL HBSS with 5 mg/mL bovine serum albumin (BSA, Sigma) with 10 μg/mL wheat-germ agglutinin (WGA) conjugated to Alexa 488 (ThermoFisher W11261), and a 1:50 dilution of Phalloidin conjugated to Alexa 405. Cells were diluted into 4.5 mL HBSS and centrifuged at 600g for 5 minutes, and resuspended in 4.5 mL HBSS and centrifuged again to wash before being resuspended in 100 μL HBSS. Cells were then plated onto LabTech II 8-well glass bottom chambers modified with 0.1% Poly-L-Lysine (PLL, Sigma P8920) and allowed to adhere for at least 4 hours at 4C before performing confocal microscopy.

#### Confocal Microscopy imaging

Confocal microscopy was performed on a Zeiss AxioVert 200M inverted microscope stand with Yokogawa CSU-22 spinning disk confocal scan head with Andor Borealis multi-point confocal system. Probes were excited by 4 laser lines in the Andor / Spectral applied Research Integrated Laser Engine: 405 nm 100 mW OPSL, 488 nm 150 mW OPSL, 561 nm 100 mW OPSL, and 642 nm 110 mW OPSL. Multipass dichroic mirror 405/488/568/647 and emission filters 450/50 nm, 525/50 nm, 605/70 nm, and 700/75 nm were used for each emission channel, respectively. Sample was imaged through a 63x oil Plan Apochromat objective with an effective pixel size of 0.092 μm/pixel. Images were captured through a Hamamatsu Orca-ER cooled CCD, and instrumentation was controlled through MetaMorph software. For each image, 9 z-planes having separation of 1.5 μm were acquired between the top and bottom of the cell, and approximately 10 fields of view were acquired for each sample.

#### Image analysis

16-bit images were read into MATLAB and converted to double precision. For each field of view, a maximum intensity projection (MIP) was calculated for the phalloidin channel. This was then binarized using adaptive thresholding, cleaned of stray pixels, and then morphological opening and closing was performed. Holes within this binarization were then filled, and discreet objects within this binarization were labeled as individual cells. For each cell in a field of view, z-planes were binarized as above using the phalloidin channel, and these z-plane binarizations were restricted to the limit of the MIP binarization for each cell. The convex hull of this z-plane binarization was used to estimate the extent of the cell, and the cell surface was estimated by selecting the perimeter of the z-plane binarization and dilating this 25 times in a 4-connected neighborhood and subsequent restriction by the undilated cell extent binarization. The total intensity of cellular probes and the surface intensity of cellular probes was calculated through summation using all z-stacks after logical indexing of the background-subtracted raw z-plane images, where background was estimated to be a constant through all z-planes and channels. Pixel-based correlation was performed through pairwise linear correlation of pixel values between channels following logical indexing. Average intensity values shown are an average over cells, and errorbars shown are the standard errors of the means, given by the standard deviation divided by sqrt(N_cells_). Internalized fraction of probe intensity for a single cell is given by (total cell intensity – cell surface intensity) / total cell intensity.

