## Supplementary Information for "Role of nanoscale antigen organization on B-cell activation probed using DNA origami"

**Supplementary Materials**
Supplementary Figs. 1–20
Supplementary Tables 1–3
Supplementary References

**Supplementary table 1. List of primers used for amplification of DNA-NP scaffolds.**

| Structure | 5'-primer (forward) | 3'-primer (reverse) |
|---|---|---|
| 6-HB | CCCTTTAGGGTTCCGATTTA | GCTGAAAAGGTGGCATCAAT |
| Icosahedron | TCTTTGCCTTGCCTGTATGA | GCTAACGAGCGTCTTTCCA |

### Supplementary Figures

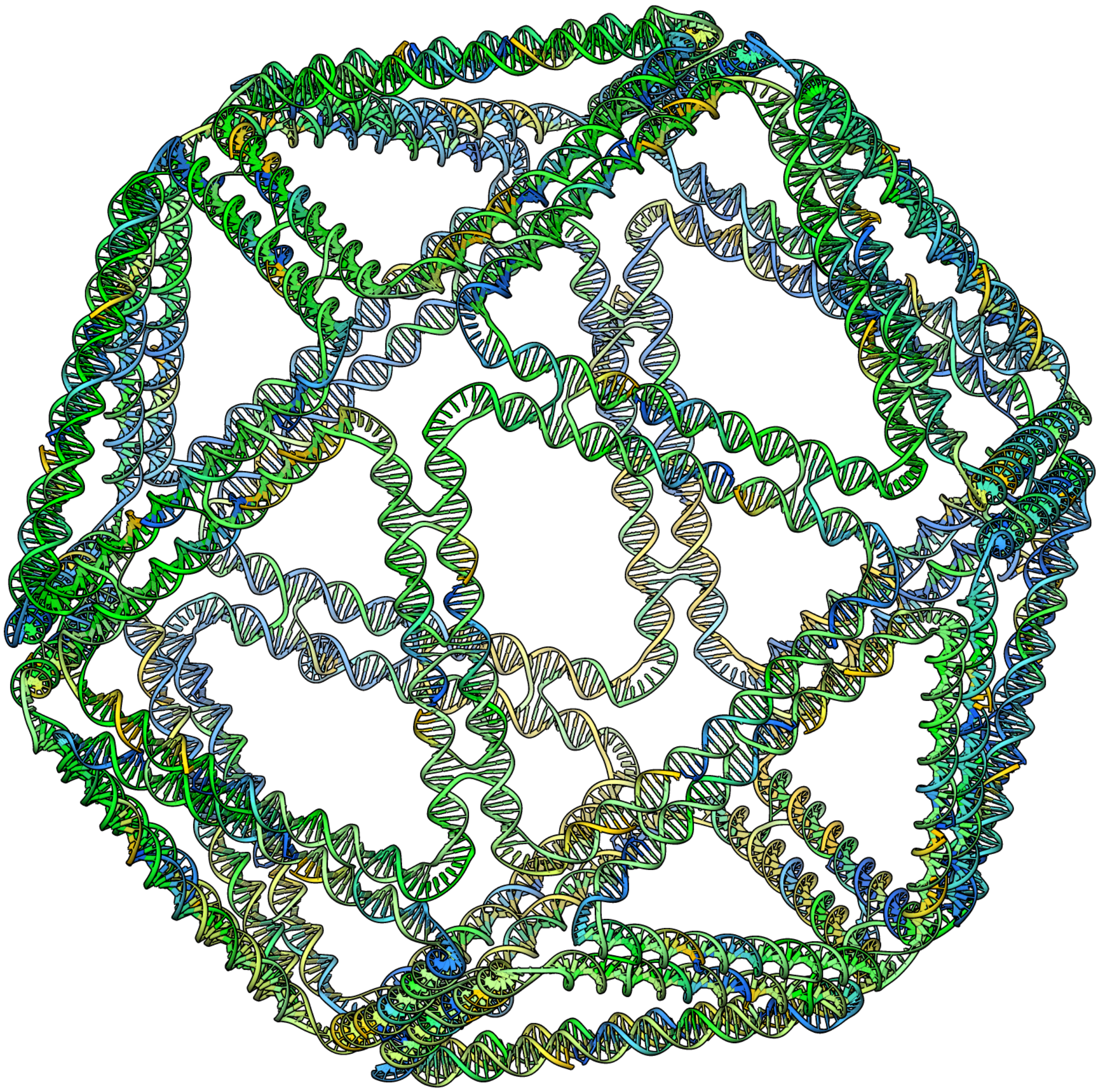

**Supplementary figure 1. All-atom model of the icosahedral scaffolded DNA origami NP.** The all-atom model was generated using DAEDALUS[1].

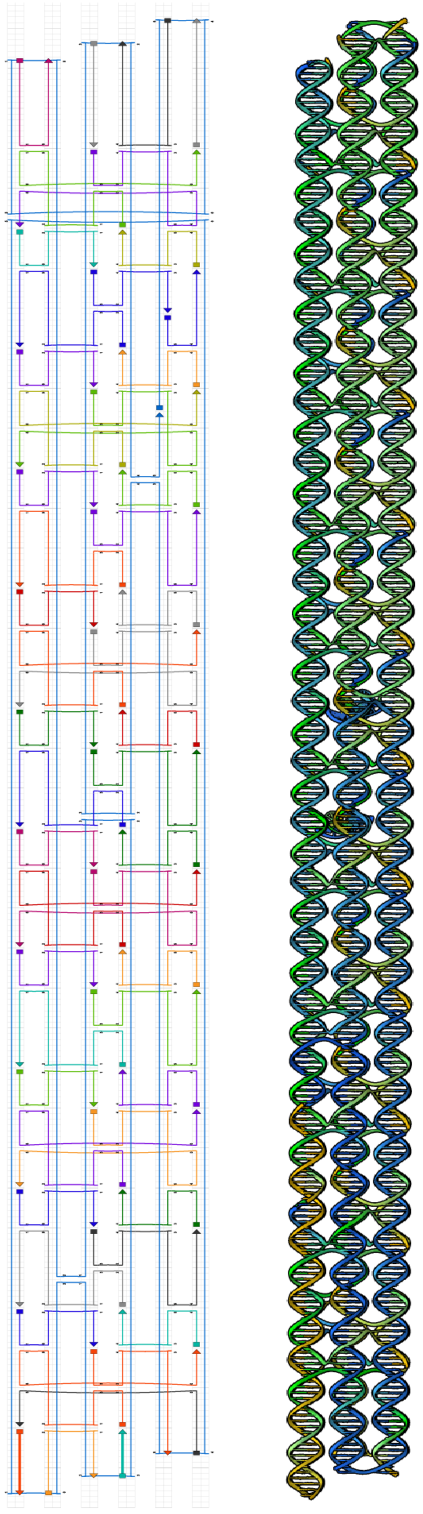

**Supplementary figure 2. Sequence design and all-atom model of the 6HB scaffolded DNA origami NP.** (*Left*) Secondary structure of the 6HB rendered using caDNAno. (*Right*) All-atom model of the 6HB DNA-NP rendered using CanDo[2,3].

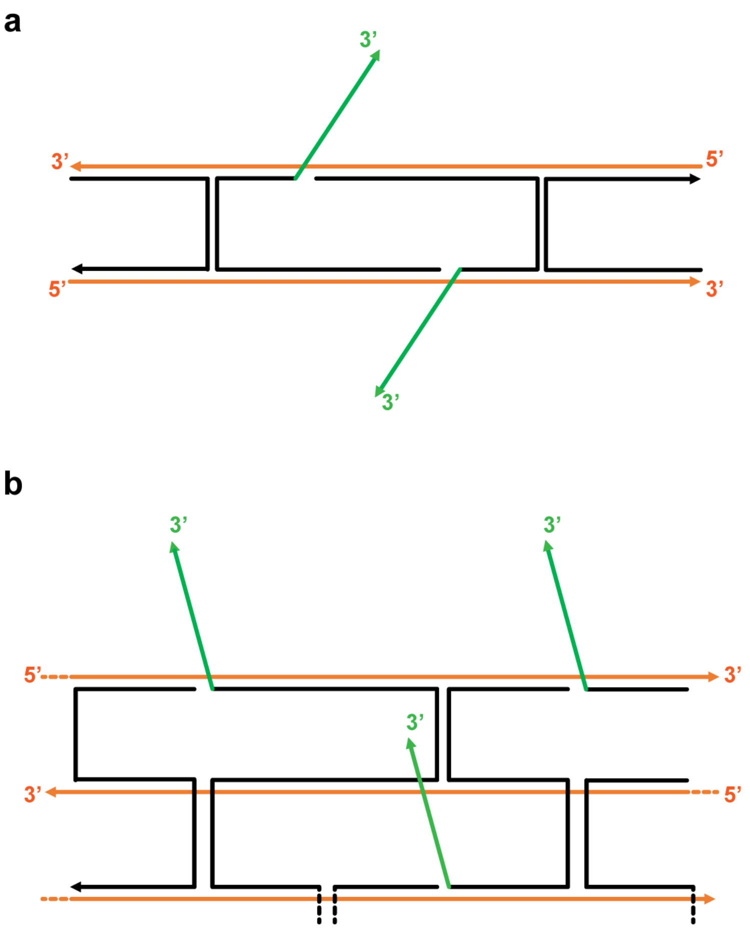


**Supplementary figure 3. Overhang placement on the edges of the DNA-NPs. (a) Secondary structure** of an edge of the DNA icosahedron with ssDNA overhangs**. (b)** Zoom-in of the secondary structure of the 6HB showing available sites for ssDNA overhangs**.**

**Supplementary Table 2. Properties of the flexible linkers used for testing the role of scaffold rigidity in antigen presentation.** Flory Radius was determined from Ma et al.,[4] and Radius-of-Gyration for the PEG polymer from Linegar et al.,[5] ($Rg=0.0215xMw^{0.583}$).

| **Linker material** | **Linker name** | **MW [Da]** | **Number of units (Bases or PEG monomers)** | **Contour Length [nm]** | **Flory Radius [nm]** | **Radius-of-Gyration [nm]** |
|---|---|---|---|---|---|---|
| ssDNA | ssDNA5 | 8930.9 | 5 | 3.2 | N/A | N/A |
| ssDNA | ssDNA12 | 11123.3 | 12 | 7.6 | N/A | N/A |
| ssDNA | ssDNA24 | 14881.8 | 24 | 15 | N/A | N/A |
| ssDNA | ssDNA35 | 18210.9 | 35 | 22 | N/A | N/A |
| ssDNA | ssDNA47 | 21916.2 | 47 | 30 | N/A | N/A |
| ssDNA | ssDNA83 | 30767.9 | 83 | 52 | N/A | N/A |
| PEG | Bis-Mal-PEG-1 | 308.3 | 2 | 0.6 | 0.5 | 0.61 |
| PEG | Bis-Mal-PEG-2 | 2000 | 45 | 12.6 | 2.8 | 1.8 |
| PEG | Bis-Mal-PEG-3 | 7500 | 170 | 47.7 | 6.1 | 3.9 |

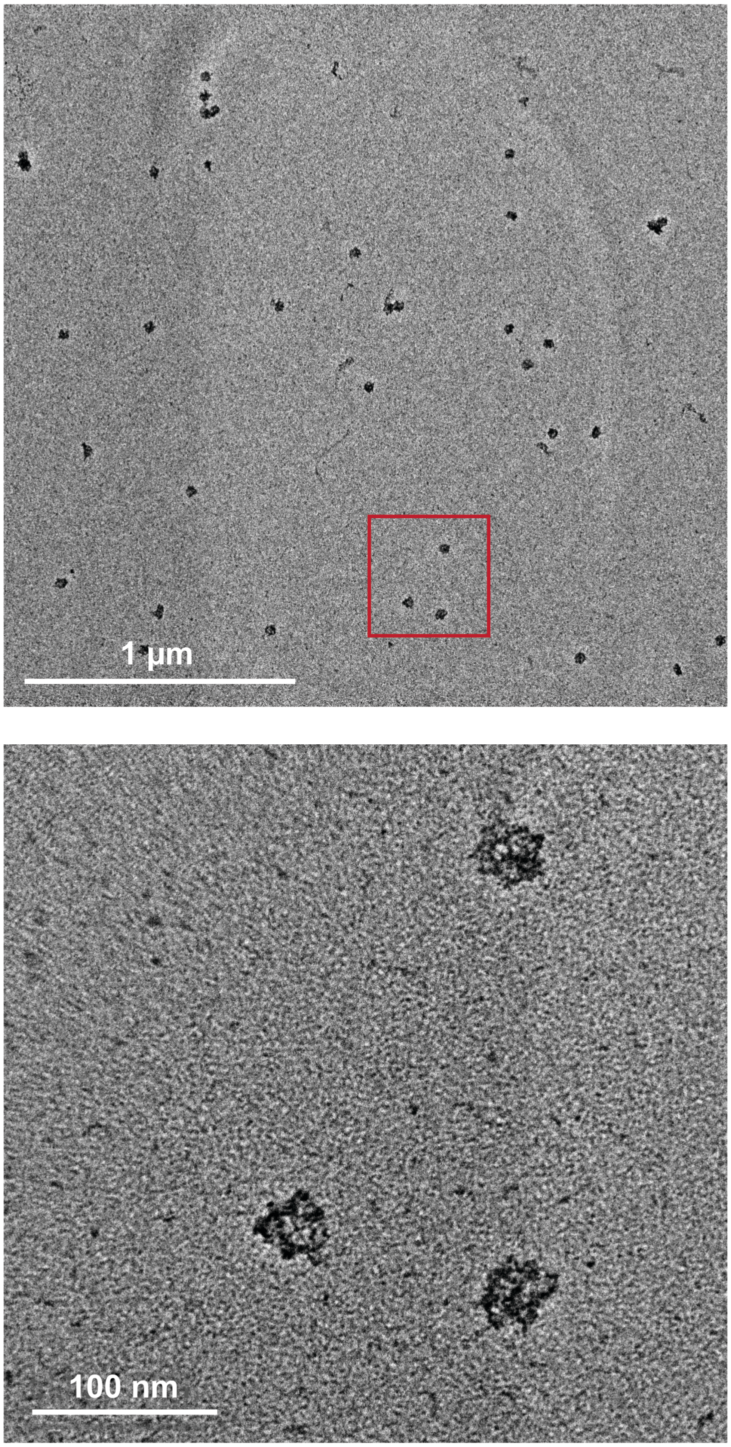


**Supplementary figure 4. TEM images of the icosahedral DNA-NP with 60 overhangs without eOD.**

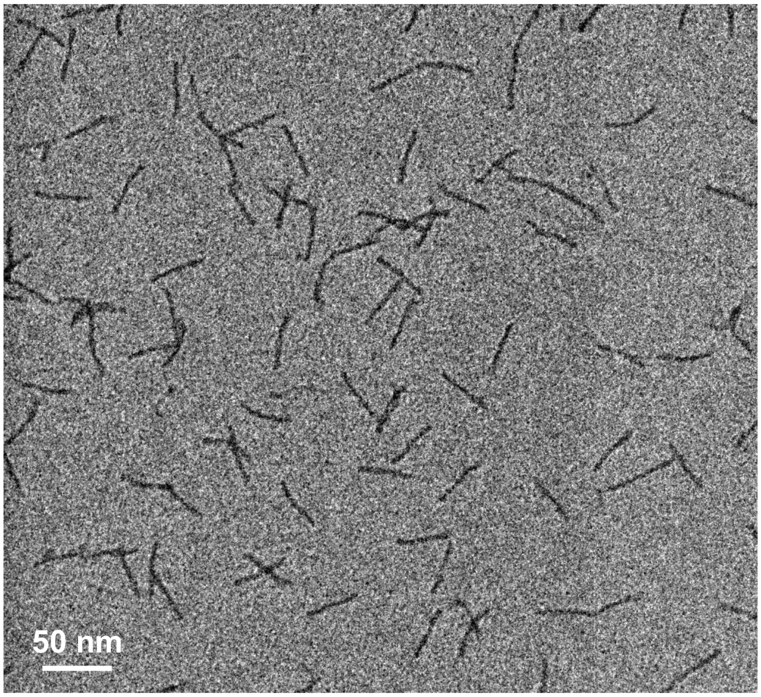


**Supplementary figure 5. TEM images of the 6HB DNA-NP with 5 overhangs without eOD.**

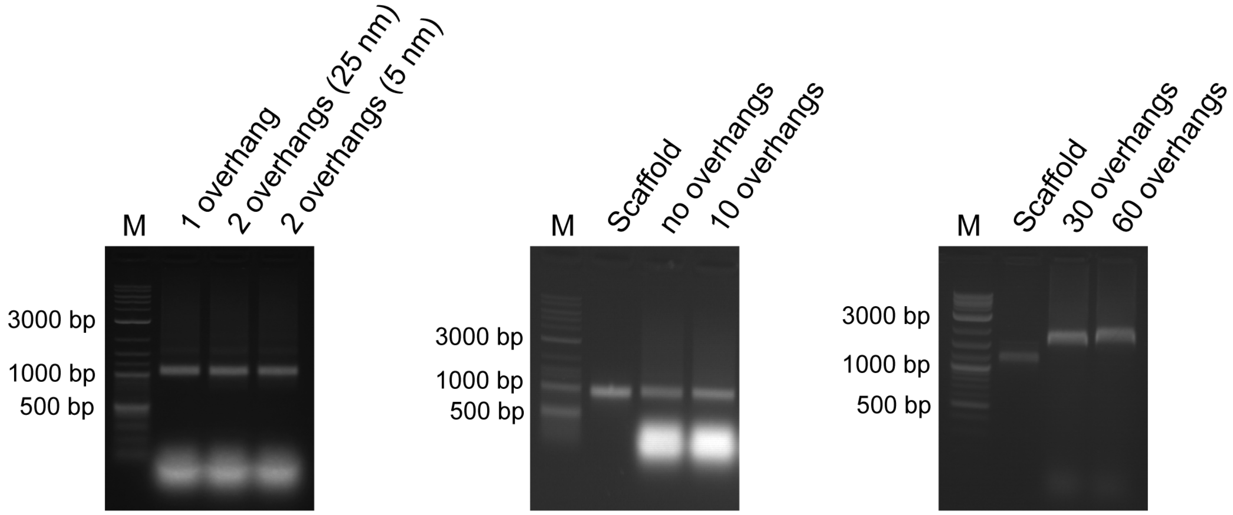


**Supplementary figure 6. Agarose gel electrophoresis of the icosahedral DNA-NP with varying numbers of overhangs.**

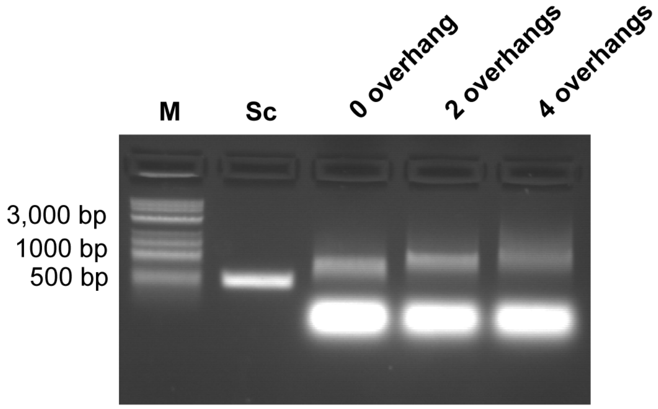


**Supplementary figure 7. Agarose gel electrophoresis of the 6HB DNA-NP with varying numbers of overhangs.**

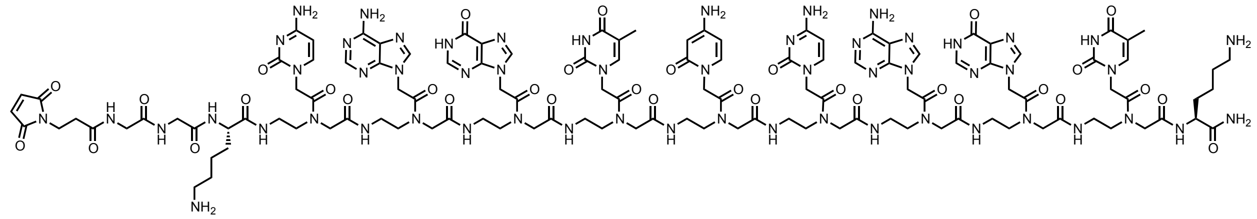

**Supplementary figure 8. Schematic of the PNA linker designed for antigen attachment to DNA-NPs.** The sequence is the following: (Maleimide)-GGK-cagtccagt-K-($CONH_2$).

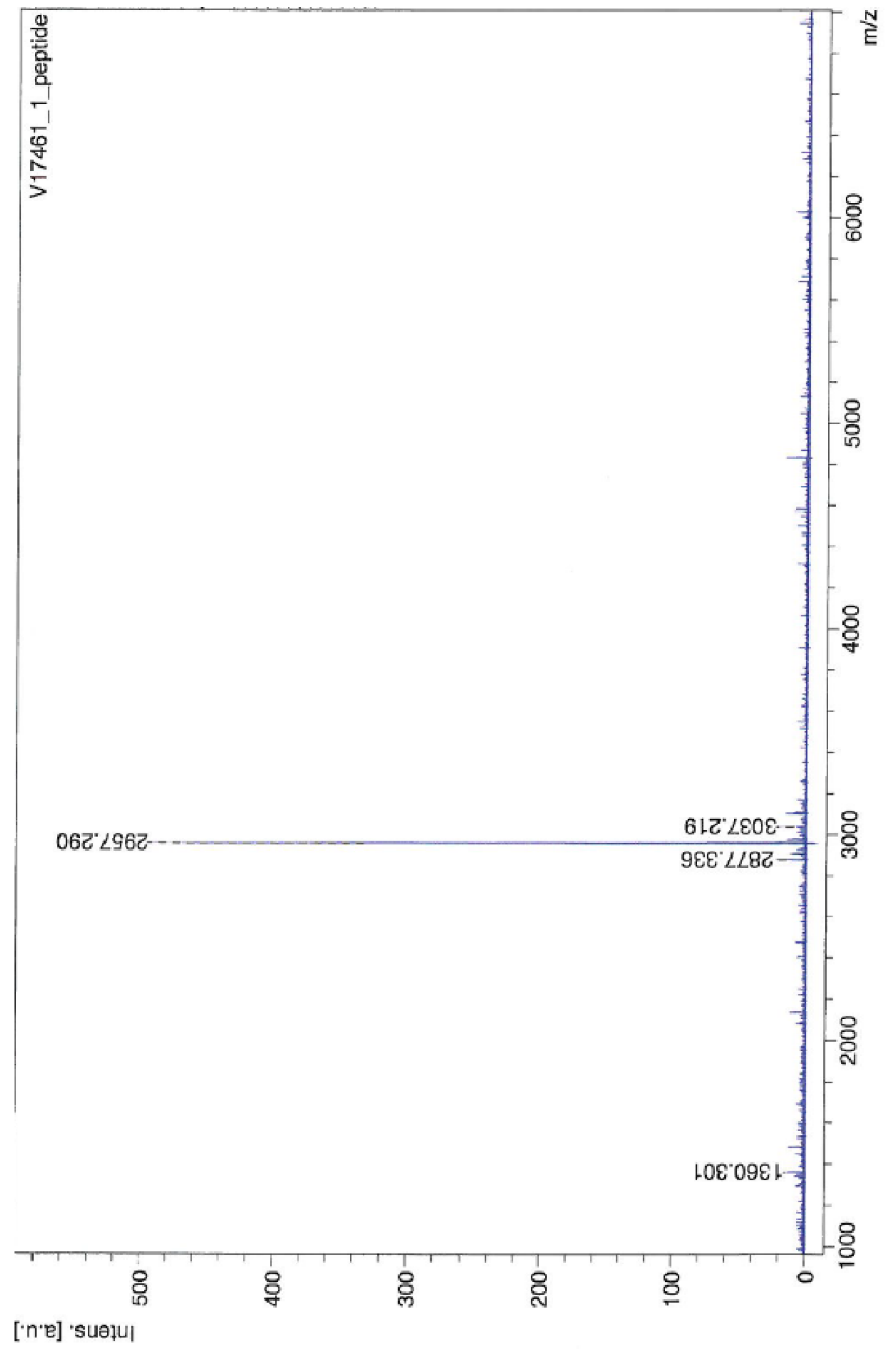


**Supplementary figure 9. MALDI-TOF analysis of the synthesized PNA strand.** Mass spectrometry analysis of maleimide modified PNA strand (expected: 2927.22 m/z, measured:

2957.29 m/z).

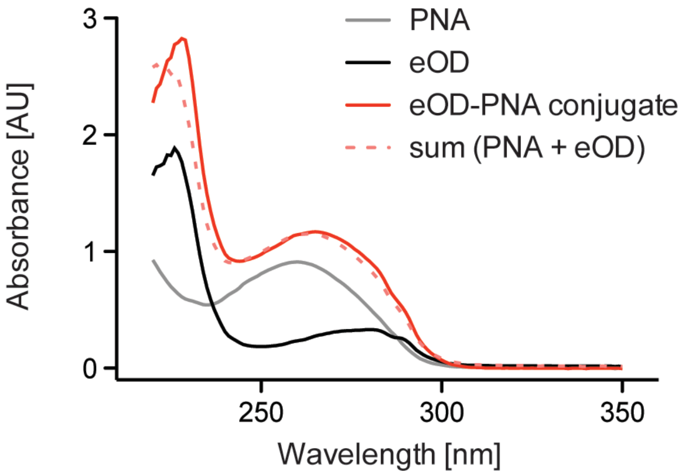


**Supplementary figure 10. Absorption spectrum analysis of PNA strand conjugation to the eOD-GT8.** UV-vis spectrum of the PNA strand alone (grey), eOD alone (black), the eOD-PNA conjugate (red), and the sum of absorbance from eOD alone and PNA alone (dotted pink).

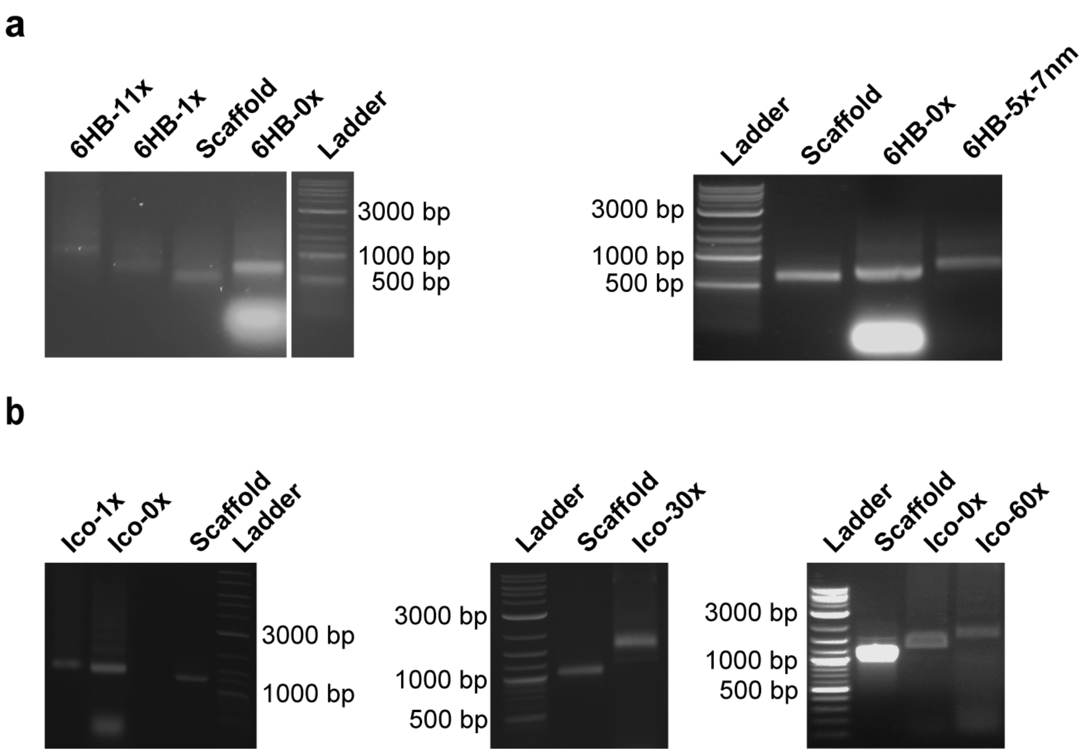


**Supplementary figure 11. eOD-GT8 antigen conjugation to DNA-NPs with ssDNA overhangs characterized with agarose gel electrophoresis.** **(a)** eOD-GT8 conjugation to various 6HB constructs. (**b**) eOD-GT8 conjugation to various icosahedral constructs. All purified structures were run on a 1.5 % agarose gel pre-stained with EtBr and run for 2 h at 70 V.

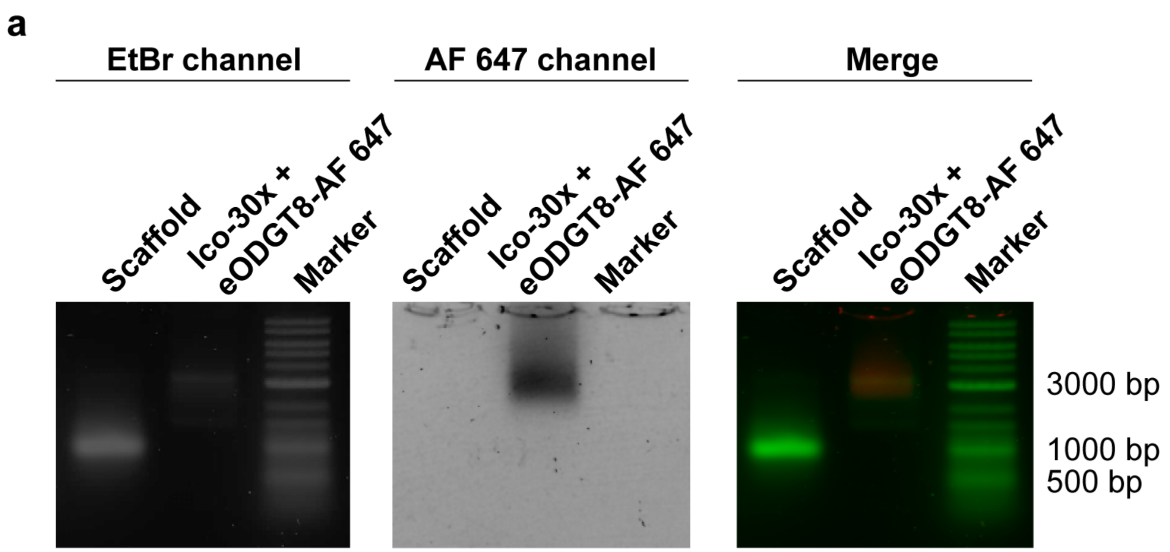


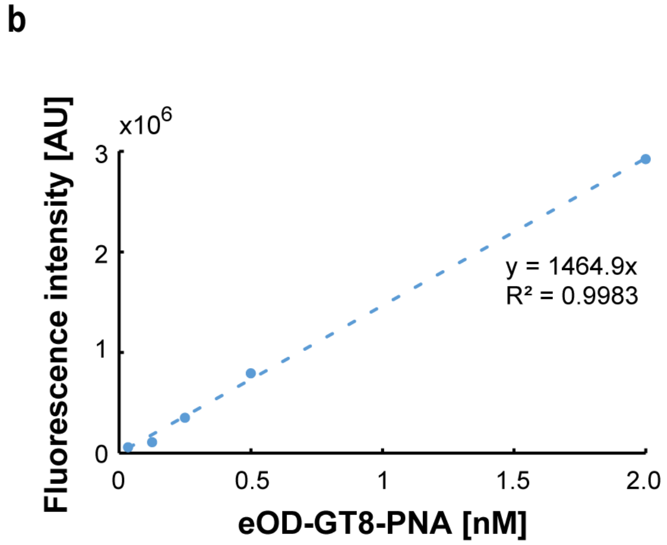


c

| Construct | Coverage [%] |
|---|---|
| Ico-1x | 107.1 ± 8.8 |
| Ico-5x | 102.8 ± 11.8 |
| Ico-30x | 89.8 ± 24.2 |
| Ico-60x | 77.3 ± 11.3 |
| 6HB-1x | 103.2 |
| 6HB-2x-7nm | 94.7 |
| 6HB-5x-7nm | 104.9 ± 18.4 |

**Supplementary figure 12. Percentage of antigen modification of the DNA-NPs determined by fluorimetry of eOD-GT8-PNA modified with AF647. a.** Fluorescent imaging of agarose gel acquired with Typhoon FLA 7000. **b**. Example of standard curve acquired with fluorescent eOD-GT8. **c**. Quantification of eOD-GT8 coverage on different DNA NPs from fluorescence spectroscopy. Error represents standard deviation of the mean (n=3 samples/group for Ico-1x, Ico-5x, Ico-30x, and 6HB-5x-7nm; n=2 samples/group for Ico-60x; n=1 samples/group for 6HB-1x, and 6HB-2x-7nm).

**a**

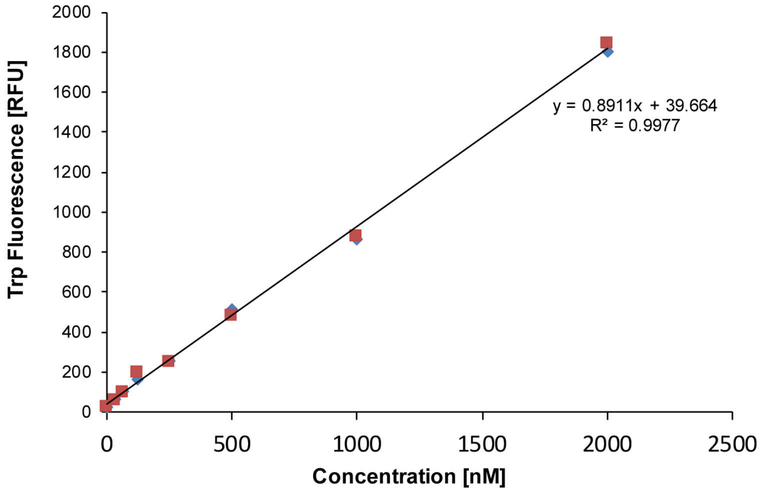


**b**

| Construct | Coverage [%] |
|---|---|
| Ico-2x | 102 |
| Ico-3x | 111 |
| Ico-4x | 102 |
| Ico-5x | 109 |
| Ico-10x | 106 |
| Ico-30x | 71.1 |

**Supplementary figure 13. Tryptophan assay to determine percentage coverage of DNA-NPs with unlabeled eOD-GT8-PNA.** (n=1 sample/groups)

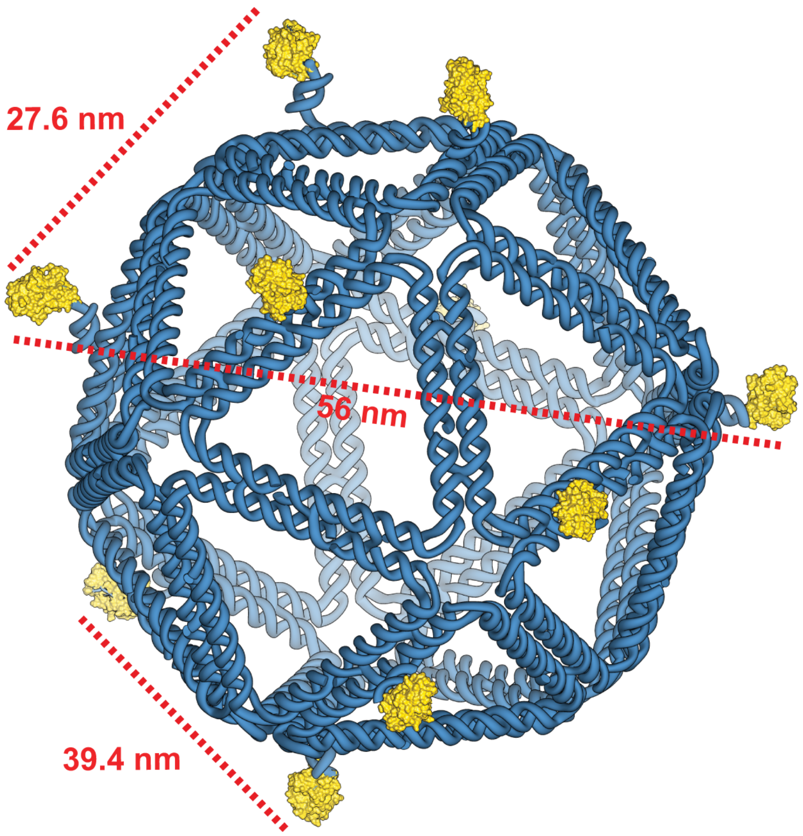


**Supplementary figure 14: Representative distances between eOD-GT8 immunogens on the icosahedral DNA-NP.** Distances were measured from the center of the antigen using UCSF Chimera[6] and are listed in Table S3.

**Supplementary Table S3. Distances between eOD-GT8 antigens in a 1, 2, 3, 4, and 5-mer icosahedral DNA-NP.** Distance were measured using UCSF Chimera[6] as shown in **Supplementary figure 14** from the center of the antigens. The error is determined using the estimated linker size (~3 nm).

| Antigen # | 1 | 2 | 3 | 4 | 5 |
|---|---|---|---|---|---|
| 1 | | 56 ± 6 nm | 43.3 ± 6 nm | 39.4 ± 6 nm | 36.5 ± 6 nm |
| 2 | 56 ± 6 nm | | 40.3 ± 6 nm | 40.9 ± 6 nm | 44.5 ± 6 nm |
| 3 | 43.3 ± 6 nm | 40.3 ± 6 nm | | 33.9 ± 6 nm | 27.6 ± 6 nm |
| 4 | 39.4 ± 6 nm | 40.9 ± 6 nm | 33.9 ± 6 nm | | 49.2 ± 6 nm |
| 5 | 36.5 ± 6 nm | 44.5 ± 6 nm | 27.6 ± 6 nm | 49.2 ± 6 nm | |

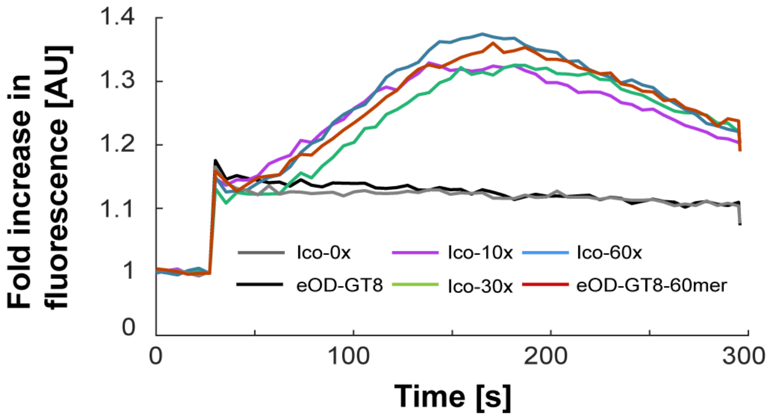


**Supplementary figure 15. B cell fluorescence response with high valency eOD-GT8 loading on the icosahedral DNA-NP.** Icosahedral DNA-NPs modified with eOD-GT8 were added to B cells loaded with Fluo-4 DNA and normalized to baseline levels. Ico-0x with 0 copy number, Ico-10x, Ico-30x, and Ico-60x contained 10, 30, and 60 copies of eOD-GT8, respectively. eOD-GT8-60mer is the positive control protein NP containing 60 copies of eOD-GT8 per NP[7] (representative individual calcium trace).

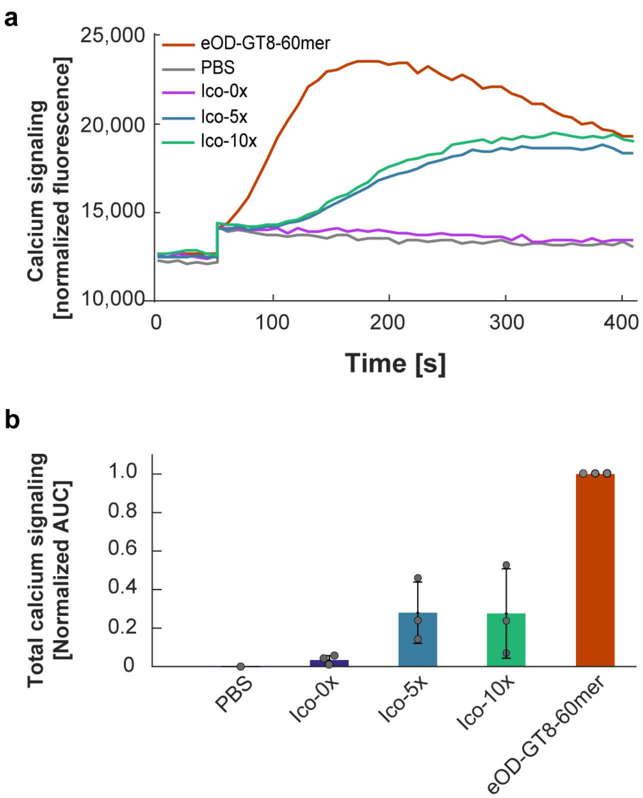


**Supplementary figure 16. Increasing valency of the low affinity antigen eOD-GT5 does not enhance B cell activation. (a)** Raw fluorescence of cells loaded with Fluo-4 calcium probe (representative individual calcium trace). (**b)** Normalized Area Under the Curve [AUC] of calcium release from cells stimulated with the icosahedral DNA-NPs displaying different valencies of the low affinity immunogen eOD-GT5. Ico-0x and Ico-10x were used at 25 nM equivalent eOD-GT8 concentration and eOD-GT8-60mer control at 2 nM equivalent eOD-GT8 concentration. Error bars represent standard deviation of the mean (n=3 samples/group).

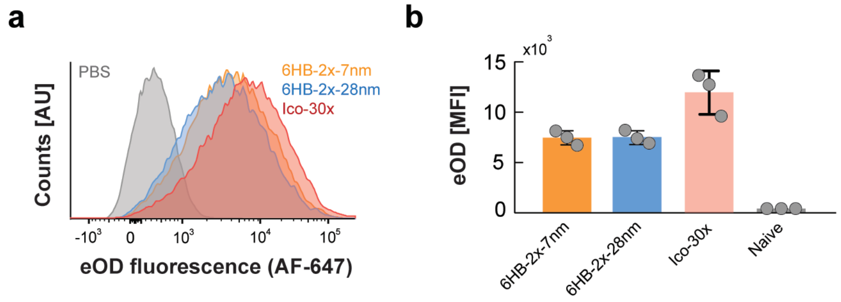


**Supplementary figure 17. Flow cytometry of labeled eOD-DNA-NPs binding to B cells.** Fluorescently labeled eOD-GT8-PNA attached to 6HB-2x-7nm, 6HB-2x-28nm, (Data as presented in **Figure 3b)** or the Ico-30x was incubated with gLVRC01 B cells for 30 minutes on ice. (**a**) Representative flow cytometry plots of Ico-30x, 6HB-2x-7nm, and 6HB-2x-28nm binding to antigen-specific B cells. (**b**) Quantitation of data from (**a**) (n = 3 samples/group).

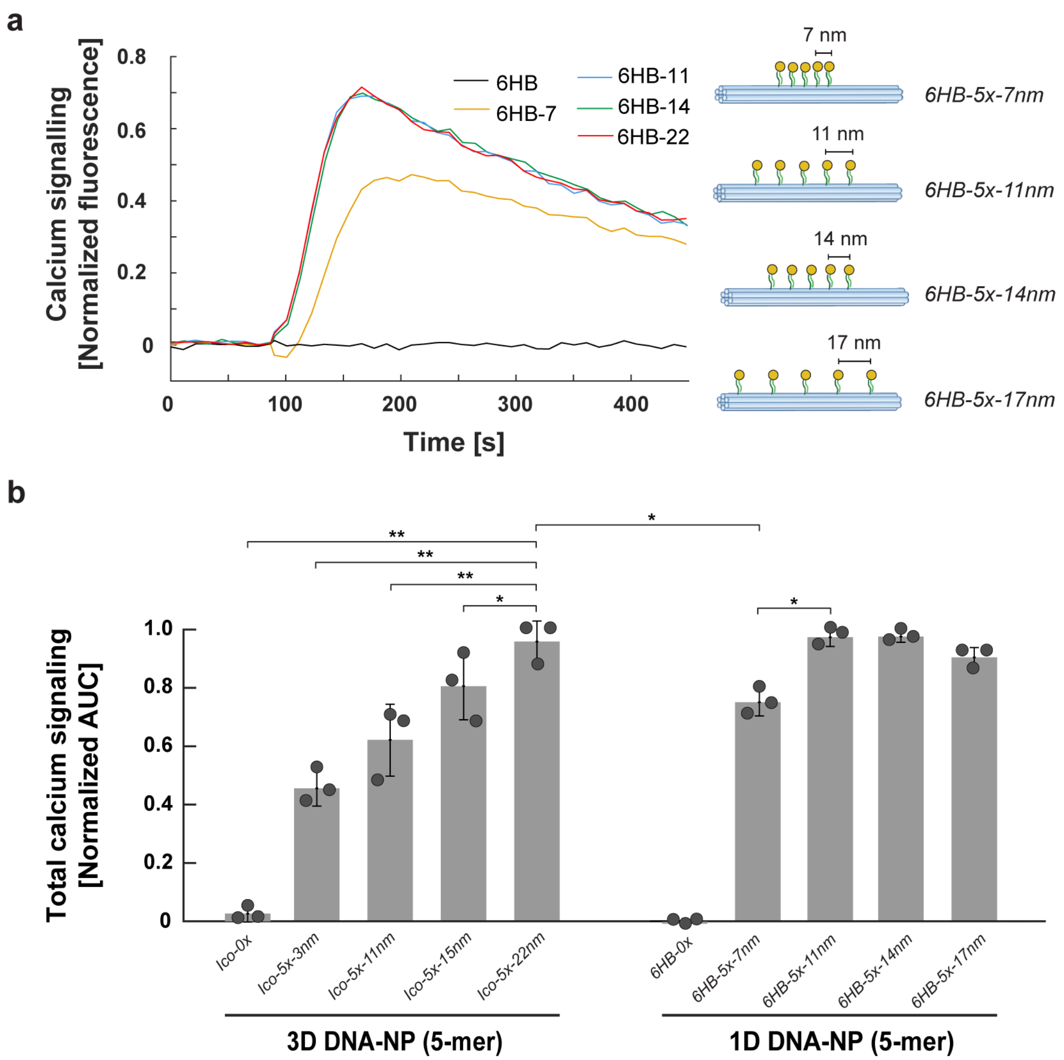


**Supplementary figure 18: Comparison of linear versus 2D clustering of five copies of antigens on Ico and 6HB. (a)** Calcium signaling of cells loaded with Fluo-4 calcium probe normalized by unstimulated levels with buffer-only control curves subtracted for 6HB structures with varying inter-antigen distances (representative individual calcium trace). (**b)** Total calcium signaling from cells stimulated with 6HB 5-mer structures presenting eOD-GT8 antigen compared with Ico 5-mer structures presented in **Fig. 4**. Fluo-4 AUC is normalized as in **Fig. 2** (n=3 samples/group), where error bars represent standard deviations and P-values are from Student's t-test (*: p<0.05; **: p<0.01; ***: p<0.001).

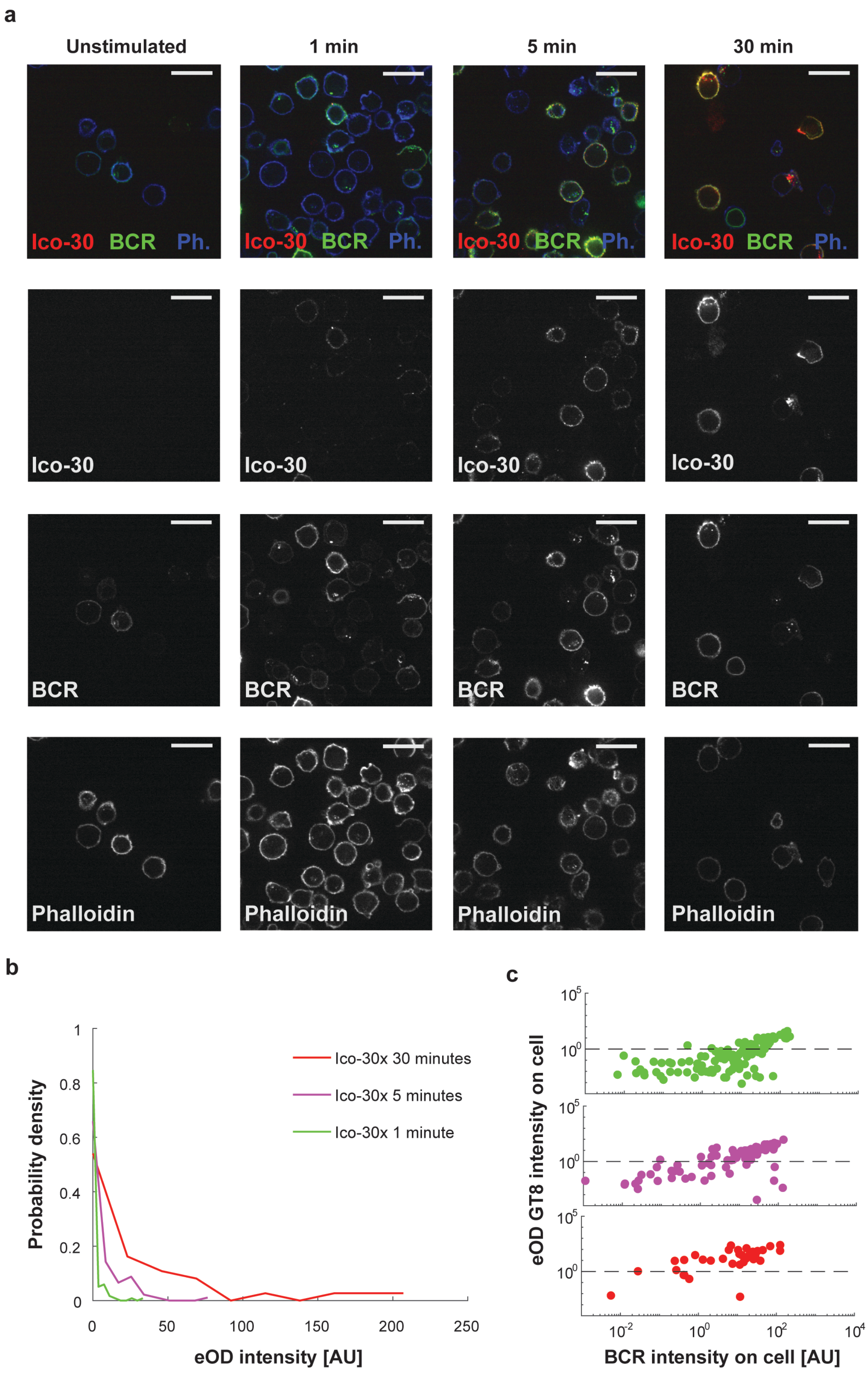
a
Unstimulated
1 min
5 min
30 min
Ico-30 BCR Ph.
Ico-30
BCR
Phalloidin
b
Probability density
eOD intensity [AU]
Ico-30x 30 minutes
Ico-30x 5 minutes
Ico-30x 1 minute
c
eOD GT8 intensity on cell
BCR intensity on cell [AU]

**Supplementary figure 19. Confocal fluorescence imaging of fixed Ramos cells with 30-mer icosahedral DNA-NP bearing fluorescent eOD.** 30-mer fluorescent eOD GT8 conjugated icosahedral DNA-NP were added to VRC01+ Ramos cells at a concentration of 5 nM and fixed in solution at various timepoints. (**a)** Images of Ramos cells with the Icosahedral 30-mer (Ico-30) bound, IgM (BCR) stained with an f(Ab)$_1$ fragment conjugated to JF549 and also stained with phalloidin (Ph.) conjugated to Alexa Fluor 405. Heterogeneity in the surface expression of the B cell receptor between B cells is apparent from the image. The binding of the icosahedral DNA-NP with fluorescent eOD is restricted to Ramos cells expressing VRC01 IgM. (**b)** The probability distribution of total eOD intensity on cells illustrates that a proportion of Ramos cells bind little or no eOD even at long times (30 minutes) following antigen addition. (**c)** Plots of individual cell total BCR intensity vs total eOD intensity further illustrate the strong correlation between surface BCR expression and eOD binding, as well as the increase in eOD binding over time. (Number of cells per conditions; 1min: 119 cells; 5 min: 103 cells; 30 min: 30 cells, from the same sample)

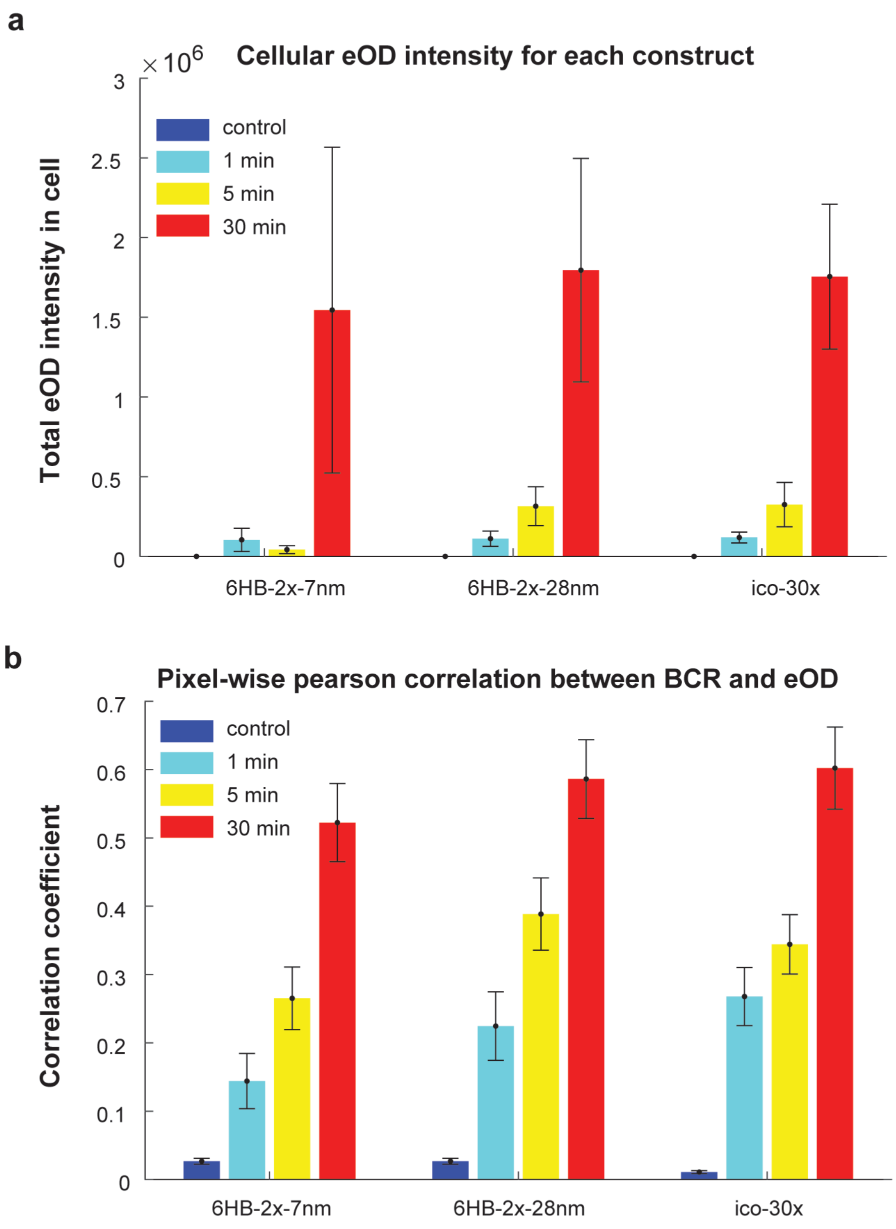


**Supplementary figure 20: Fluorescence quantification of confocal microscopy images.** Fluorescent eOD GT8 conjugated Ico-30x was added to VRC01+ Ramos cells at a concentration of 5 nM and fixed in solution at various timepoints. (**a**) Average total eOD fluorescence per cell increases over time for all constructs. (**b**) The pixel-based Pearson correlation between eOD and BCR intensity also increases over time. Total fluorescence is calculated across all z-sections and average is computed across all cells. Error bars represent standard errors of the means across all cells per condition. (Number of cells per conditions; **Ico-30x**: 1min: 14 cells, 5 min: 183 cells, 30 min: 15 cells; **6HB-2x-7nm**: 1min: 26 cells, 5 min: 22 cells, 30 min: 19 cells; **6HB-2x-28nm**: 1min: 15 cells, 5 min: 26 cells, 30 min: 23 cells; **Control**: 29 cells)
